# Transcription regulates bleb formation and stability independent of nuclear rigidity

**DOI:** 10.1101/2022.11.14.516344

**Authors:** Isabel K. Berg, Marilena L. Currey, Sarthak Gupta, Yasmin Berrada, Bao Nyugen Viet, Mai Pho, Alison E. Patteson, J. M. Schwarz, Edward J. Banigan, Andrew D. Stephens

## Abstract

Chromatin is an essential component of nuclear mechanical response and shape that maintains nuclear compartmentalization and function. The biophysical properties of chromatin alter nuclear shape and stability, but little is known about whether or how major genomic functions can impact the integrity of the nucleus. We hypothesized that transcription might affect cell nuclear shape and rupture through its effects on chromatin structure and dynamics. To test this idea, we inhibited transcription with the RNA polymerase II inhibitor alpha-amanitin in wild type cells and perturbed cells that present increased nuclear blebbing. Transcription inhibition suppresses nuclear blebbing for several cell types, nuclear perturbations, and transcription inhibitors. Furthermore, transcription is necessary for robust nuclear bleb formation, bleb stabilization, and bleb-based nuclear ruptures. These morphological effects appear to occur through a novel biophysical pathway, since transcription does not alter either chromatin histone modification state or nuclear rigidity, which typically control nuclear blebbing. We find that active/phosphorylated RNA pol II Ser5, marking transcription initiation, is enriched in nuclear blebs relative to DNA. Thus, transcription initiation is a hallmark of nuclear blebs. Polymer simulations suggest that motor activity within chromatin, such as that of RNA pol II, can generate active forces that deform the nuclear periphery, and that nuclear deformations depend on motor dynamics. Our data provide evidence that the genomic function of transcription impacts nuclear shape stability, and suggests a novel mechanism, separate and distinct from chromatin rigidity, for regulating large-scale nuclear shape and function.

## Introduction

Aberrant alterations in transcription and disruptions to nuclear shape are two common cellular hallmarks of human disease. Both are used as prognostic indicators of disease severity in breast, cervical, and prostate cancers, among others (Papanicolaou and Traut, 1997; Helfand *et al*., 2012; Radhakrishnan *et al*., 2017; Lu *et al*., 2018). Several studies have suggested that there are interactions between transcription and nuclear shape. In the testosterone-sensitive prostate cancer model cell line LNCaP, testosterone-induced transcription via the androgen receptor results in increased nuclear blebbing (Helfand *et al*., 2012). Transcription activation via TGFβ1 has also been shown to induce abnormal nuclear shape (Chi *et al*., 2022). Additionally, active RNA polymerase II and transcriptionally active genes and chromosomes are enriched in nuclear blebs in cells with lamin B1 knocked down by shRNA and in progeria cells (Shimi *et al*., 2008; Bercht Pfleghaar *et al*., 2015). The biochemical and biophysical state of chromatin can control nuclear morphology (Furusawa *et al*., 2015; Schreiner *et al*., 2015; Stephens *et al*., 2018), but it is unclear if transcription is necessary to induce abnormal nuclear shape in perturbed nuclei. Furthermore, the mechanism of how transcription affects nuclear shape is unknown. Thus, understanding how transcription affects nuclear shape and integrity would provide new insights to these two vital functions that are perturbed in disease.

Chromatin mechanics is essential to nuclear mechanical response, and it is a key determinant of nuclear shape and stability (Stephens *et al*., 2019b). Chromatin generates a spring-like elastic response to small, few-microns-sized mechanical strains, complementing the robust elastic response to large deformations provided by lamins (Stephens *et al*., 2017; Hobson *et al*., 2020; Currey *et al*., 2022). The rigidity of chromatin is governed by histone modifications (Chalut *et al*., 2012; Krause *et al*., 2013; Shimamoto *et al*., 2017; Stephens *et al*., 2018, 2019a; Hobson *et al*.,2020; Nava *et al*., 2020), H1 dynamics (Furusawa *et al*., 2015; Senigagliesi *et al*., 2019), chromatin-binding proteins (Schreiner *et al*., 2015; Wang *et al*., 2018; Williams *et al*., 2020; Strom *et al*., 2021), and 3D genome organization (Belaghzal *et al*., 2021). Independent of lamins, chromatin decompaction results in a softer nucleus, which can more easily succumb to external perturbations, such as cytoskeletal forces. Thus, the nucleus can lose its normal shape and form a protrusion called a nuclear bleb (Furusawa *et al*., 2015; Stephens *et al*., 2018). Both chromatin and lamin perturbations result in nuclear blebs, which have highly curved surfaces prone to rupture (De Vos *et al*., 2011; Vargas *et al*., 2012; Stephens *et al*., 2018; Xia *et al*., 2018; Nmezi *et al*., 2019; Pfeifer *et al*., 2022). The resulting loss of nuclear compartmentalization via blebbing and rupture causes genomic dysfunction through DNA damage (Denais *et al*., 2016; Irianto *et al*.,2016; Raab *et al*., 2016; Chen *et al*., 2018; Xia *et al*., 2018; Stephens *et al*., 2019a; Shah *et al*., 2021), global changes to transcription along with transcriptional inhibition within the bleb (De Vos *et al*., 2011; Helfand *et al*., 2012), and loss of cell cycle control (Pfeifer *et al*., 2018). Thus, the physical properties of chromatin, and potentially transcription, can impact nuclear function through nuclear morphology.

Transcription affects both chromatin structure and dynamics, which suggests possible pathways to nuclear blebbing and rupture. Chromosome conformation capture (Hi-C and Micro-C) experiments have indicated that polymerases may aid in the establishment of A/B chromatin compartments typically associated with eu- and heterochromatin (Jiang *et al*., 2020; Zhang *et al*., 2021), generate characteristic spatial patterns around active genes (Banigan *et al*., 2022; Zhang *et al*., 2022), and contribute to enhancer-promoter contacts (Hsieh *et al*., 2020; Zhang *et al*., 2022). Furthermore, active transcription is generally associated with open, decompact chromatin structure, as measured by ATAC-seq (Buenrostro *et al*., 2013). However, it has also been suggested that transcription could play a restrictive role on chromatin by locally constraining it via crosslinking (Nagashima *et al*., 2019) or phase separation (Hnisz *et al*., 2017). Nonetheless, transcription drives large-scale chromatin dynamics, which manifest as correlated motions of micron-sized regions (Zidovska *et al*., 2013; Shaban *et al*., 2018, 2020). Simulations suggest that chromatin connectivity combined with polymerase motor activity could generate these dynamics (Liu *et al*., 2021). The same mechanism could contribute to nuclear shape fluctuations of isolated nuclei (Liu *et al*., 2021), although experiments indicate that transcription inhibition in live cells marginally enhances shape fluctuations (Chu *et al*., 2017). Thus, transcriptional changes could potentially affect nuclear morphology and compartmentalization through changes to chromatin-based nuclear mechanical response or large-scale chromatin dynamics.

To address transcription’s role in determining nuclear shape, mechanics, and ruptures we inhibited transcription using the RNA pol II inhibitor alpha-amanitin in wild type cells and cells treated with the histone deacetylase inhibitor (HDACi) valproic acid (VPA). In this way, we assessed the nuclear morphological effects of transcription under both normal conditions and conditions that typically result in nuclear blebs. Treatment with alpha-amanitin did not change nuclear blebbing in both mouse embryonic fibroblasts (MEFs) and HT1080 wild type cells, but it suppressed nuclear blebbing in VPA-treated cells. Micromanipulation force measurements showed that transcription inhibition did not alter nuclear rigidity, suggesting an alternative, dynamic mechanism for nuclear blebbing. Time-lapse imaging of cells with GFP-tagged nuclear localization signal (NLS-GFP) to track nuclear blebbing and ruptures revealed that transcription is essential for nuclear bleb formation and stabilization, which ultimately affects the number of nuclear ruptures that occur per nucleus. We found that active initiating RNA pol II, but not elongating RNA pol II, was enriched in nuclear blebs. These findings can be recapitulated and understood through a polymer simulation model in which chromatin is a crosslinked polymer contained within a polymer shell (the lamina), and transcription is represented by motor activity within the chromatin. Thus, we find that transcription is a novel, dynamic mechanism of nuclear blebbing and ruptures, that acts without changing the bulk rigidity of chromatin.

## Results

### Alpha-amanitin decreases transcriptional activity in wild type and VPA-treated cells

Transcription is a major nuclear function that affects the biophysical properties of chromatin and the nucleus, and thus could modulate nuclear shape and dynamics. To assess this possibility, we inhibited transcription using the RNA polymerase II inhibitor alpha-amanitin in wild type mouse embryonic fibroblast (MEF) cells that were untreated or treated with the histone deacetylase inhibitor (HDACi) valproic acid (VPA). VPA induces chromatin decompaction via increased euchromatin, which results in a weaker nucleus that blebs and ruptures (Stephens *et al*., 2018, 2019a).

To assay changes in transcription upon alpha-amanitin treatment, we measured whole nuclear RNA levels by uridine analog 5-ethynyluridine (EU) and click chemistry (Jao and Salic, 2008). VPA treatment resulted in increased RNA levels compared to untreated (unt) wild type cells (**Figure 1, A and B**). Treatment with RNA pol II inhibitor alpha-amanitin (aam) drastically reduced nuclear RNA levels in both wild type and VPA-treated cells. Thus, alpha-amanitin successfully represses transcription in both untreated and VPA-treated cells.

**Figure 1.**
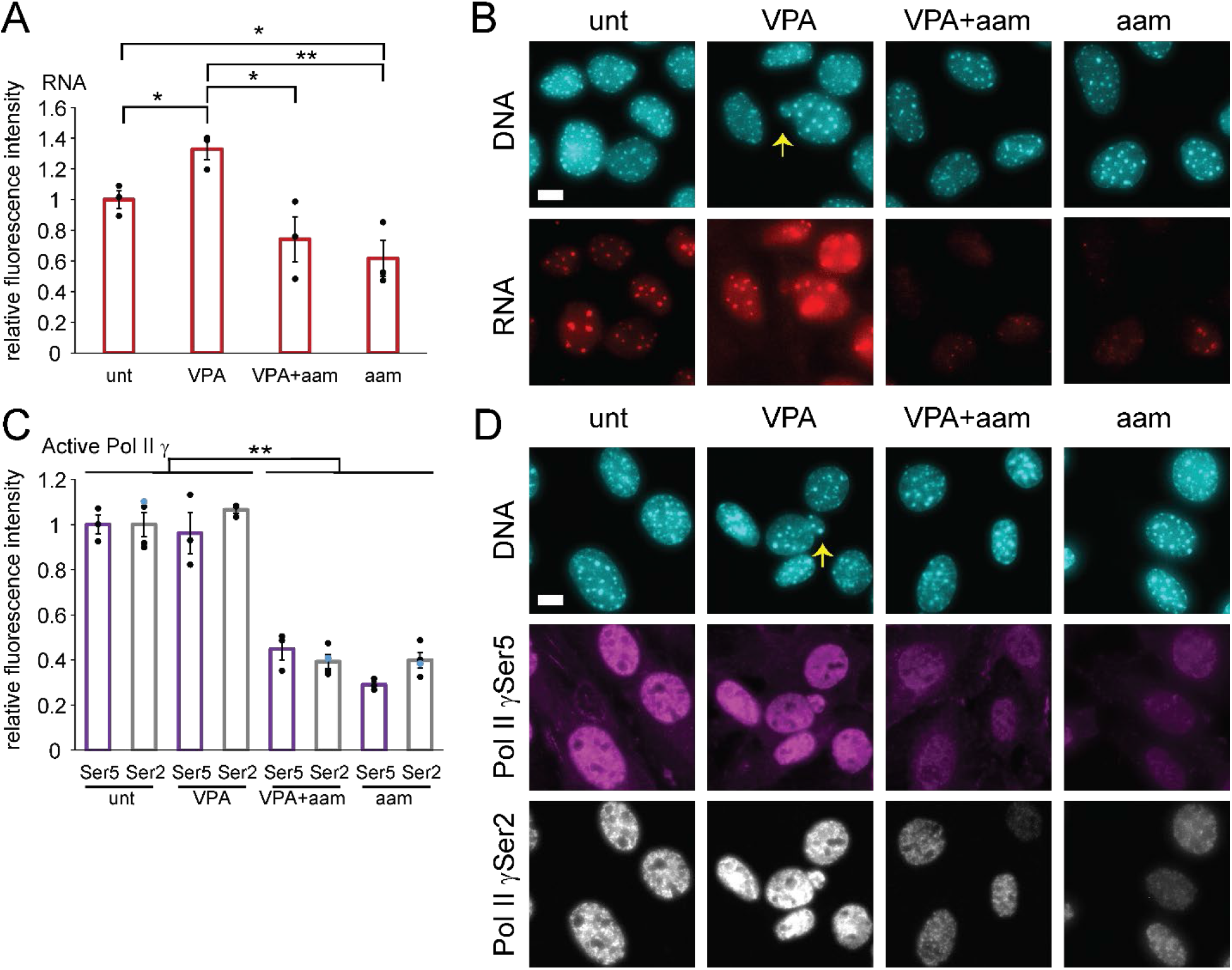
Transcription is decreased upon treatment with RNA pol II inhibitor alpha-amanitin in both untreated and VPA. (A) Graph and (B) example images of RNA immunofluorescence levels in MEFs, labeled via EU click-it-chemistry (red) and DNA labeled via Hoechst (cyan) for untreated (unt), VPA-treated (VPA), VPA- and alpha-amanitin-treated (VPA+aam), and alpha-amanitin-treated (aam) cells. (C) Graph and (D) example images of immunofluorescence levels of RNA Polymerase II phosphorylated at Ser5 (purple, initiating) and Ser2 (gray, elongating) with DNA labeled via Hoechst (cyan). Yellow arrow denotes nuclear bleb in example images. Three biological replicates with n > 30 cells. Error bars represent standard error and statistical test are Student’s t-tests, with significance denoted by * p < 0.05, ** p < 0.01, and *** p < 0.001. Scale bar is 10 μm.

Next, we aimed to establish how VPA and alpha-amanitin affect active RNA pol II levels. We measured the average immunofluorescence intensities in nuclei of active RNA pol II phosphorylated at Ser5 (initiating) and Ser2 (elongating). Without alpha-amanitin, we found no difference in levels of RNA pol II γSer5 and γSer2 in untreated cells and VPA-treated cells (**Figure 1, C and D**). As expected, alpha-amanitin treatment resulted in a drastic 60% or greater decrease of active RNA pol II in cells without or with VPA treatment (**Figure 1, C and D**). The decreased levels of active RNA pol II were also similar between γSer5 and γSer2, suggesting both initiation and elongation are inhibited equally. Thus, alpha-amanitin decreases transcription via inhibition of active RNA pol II.

### Transcription is a major component of nuclear blebbing

With a defined approach for decreasing transcription, we aimed to determine the effect that transcription has on nuclear blebbing. Untreated wild type nuclei normally exhibit a low background level of nuclear blebbing, with 4% of nuclei presenting blebs. With VPA treatment, the percentage of nuclei with nuclear blebs increased to 11% (**Figure 2A**), consistent with previous reports (Stephens *et al*., 2018, 2019a). Interestingly, dual treatment with the RNA pol II inhibitor alpha-amanitin (aam) and VPA for 16-24 hours resulted in a significant decrease in nuclear blebbing levels from 11% in VPA to 6% in VPA and alpha-amanitin, effectively returning nuclear blebbing to wild type levels (**Figure 2A**). Addition of alpha-amanitin alone does not alter wild type nuclear blebbing percentages, as they remain at 4%. Thus, transcription inhibition is a significant inhibitor of nuclear blebs in cells with chromatin decompaction induced by VPA.

**Figure 2.**
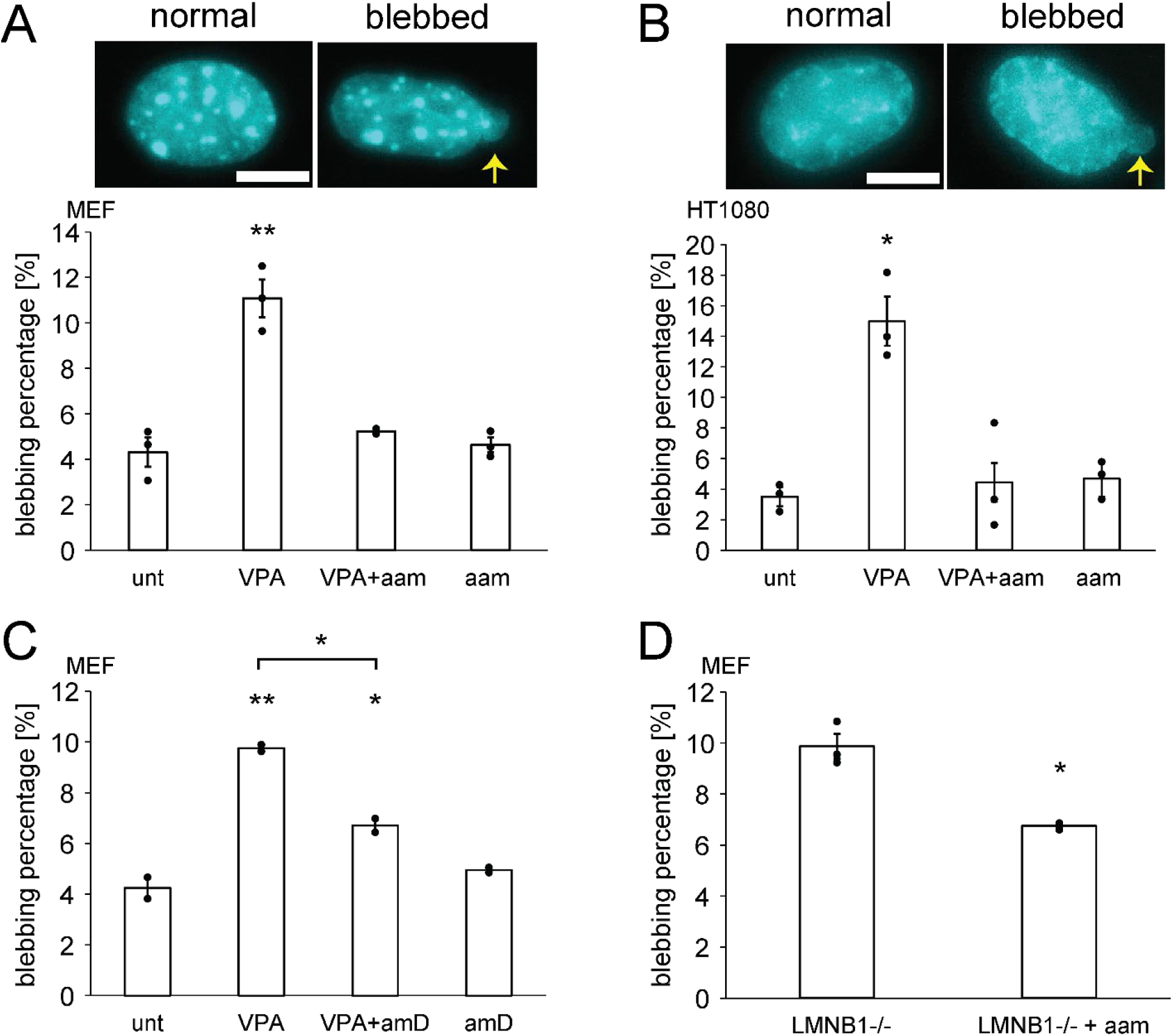
Transcription inhibition suppresses nuclear blebbing across cell types, drugs, and perturbations that cause nuclear blebbing. Example images and graph of percentages of nuclei that bleb in (A) MEF cells and (B) HT1080 cells for untreated (unt), VPA-treated (VPA), VPA- and alpha-amanitin-treated (VPA+aam), and alpha-amanitin-treated (aam) cells. Three biological replicates with n = 60-200 cells each are shown as dots. Yellow arrow denotes nuclear bleb in example images. (C) Graph of nuclear blebbing percentages in MEF cells for untreated or VPA-treated cells with or without pol II inhibitor actinomycin D (amD). Two biological replicates with n > 450 cells each are shown as dots. (D) Graph of nuclear blebbing percentages in MEF LMNB1−/− cells without or with the RNA pol II inhibitor alpha-amanitin (aam). Three biological replicates with n > 550 cells each are shown as dots. Error bars represent standard error and statistical tests are Student’s t-tests, with significance denoted by * p < 0.05, ** p < 0.01, and *** p < 0.001. Scale bar is 10 μm.

Next, we aimed to determine if the effect of transcription inhibition on nuclear blebbing is a general phenomenon. First, we considered an alternative cell type. HT1080 cells also display increased levels of nuclear blebbing upon chromatin decompaction by VPA treatment, from 3% to 15% (**Figure 2B**), as previously reported (Stephens *et al*., 2019a). Similar to MEFs, dual treatment of HT1080 cells with alpha-amanitin and VPA resulted in wild type levels of nuclear blebbing, indicating that bleb formation resulting from VPA treatment was suppressed. Next, we used an alternative transcription inhibitor, actinomycin D (amD), to treat wild type or VPA-treated MEFs. This treatment also resulted in a significant decrease in nuclear blebbing induced by VPA (**Figure 2C**). Finally, we used an alternative nuclear perturbation to induce blebbing. MEF LMNB1−/− cells, null for lamin B1, also result in a significant increase in nuclear blebbing relative to wild type cells (Lammerding *et al*., 2006; Vargas *et al*., 2012; Hatch and Hetzer, 2016; Stephens *et al*., 2019a; Young *et al*., 2020). MEF LMNB1−/− cells treated with alpha-amanitin exhibited decreased levels of nuclear blebbing, dropping significantly from 10% to 6% (**Figure 2D).** Thus, nuclear blebbing is dependent on active transcription across different cell types, transcription inhibition treatments, and perturbations that increase nuclear blebbing.

### Nuclear rigidity is not altered by transcription inhibition

Nuclear blebbing is thought to be driven by disruptions to one or both of the two major nuclear mechanical components, chromatin and lamin A/C (Kalukula *et al*., 2022). We thus investigated whether transcription inhibition can alter nuclear mechanics.

Since VPA treatment leads to softer nuclei due to increased histone acetylation, we sought to determine whether transcription inhibition suppresses bleb formation by suppressing histone acetylation. To this end, we assayed levels of euchromatin via H3K9ac immunofluorescence. With VPA treatment, euchromatin levels increased by ~50% from untreated wild type cells, as previously reported (Stephens *et al*., 2017, 2018). Surprisingly, treatment with alpha-amanitin and VPA together increased the levels of euchromatin even more than VPA treatment alone, but alpha-amanitin treatment alone did not have any effect on euchromatin levels (**Figure 3, A and B**). This data demonstrates that nuclear bleb suppression in transcriptionally inhibited VPA-treated cells does not result from decreases in euchromatin levels.

**Figure 3.**
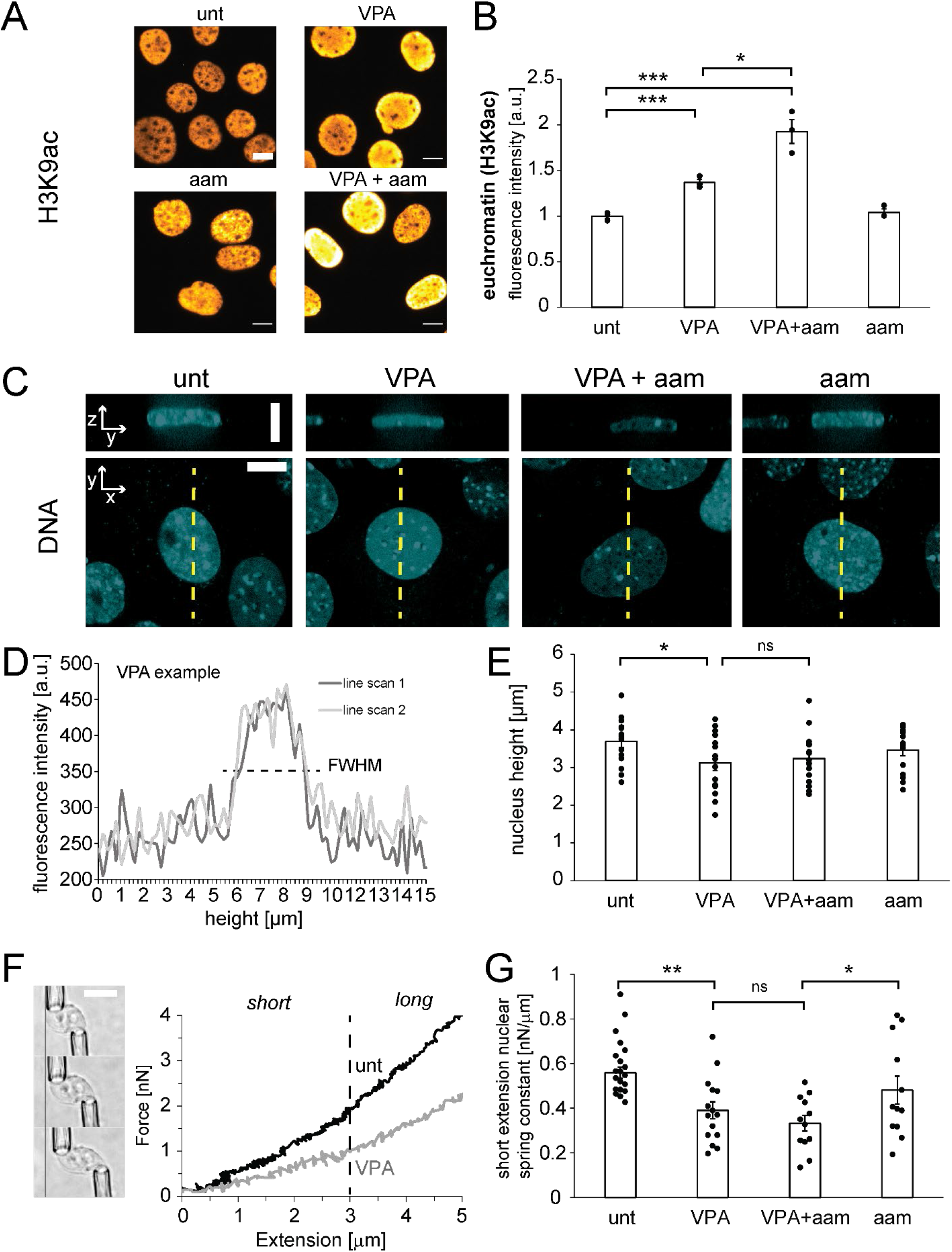
The mechanical properties of the nucleus are not altered by transcription inhibition. (A) Example images and (B) graphs of relative immunofluorescence signal of euchromatin (H3K9ac) in untreated (unt), VPA-treated (VPA), VPA- and alpha-amanitin-treated (VPA+aam), and alpha-amanitin-treated cells (aam). Three biological replicates represented as dots consisted of 100 - 300 cells each. (C) Example images of side and top-down views of Hoeschst-labeled nuclei captured using spinning disk confocal microscopy. Yellow dotted line denotes cross section. (D) Example line scans through nucleus of VPA-treated cell’s side-on image from panel C, where full-width at half maximum (FWHM) from two different line scans were averaged to determine the height of each nucleus. (E) Individual (dots) and average (bar) measurements of nuclear height for each condition (n = 15 nuclei for each condition). (F) *Left:* Example images of a micromanipulation force-extension measurement, with an isolated nucleus pulled by the “pull” pipette (bottom right) to extend the nucleus and held by a the “force” pipette (top left), whose deflection multiplied by a precalibrated spring constant measures force. *Right:* Example force-extension graph of control (unt) and chromatin-decompacted (VPA) cells over short and long regimes. (G) Graph of the individual (dots) and average (bar) short-extension nuclear spring constants (unt, n = 21; VPA, n = 15; VPA+aam, n = 12; aam, n = 12). Long-extension spring constants do not change for all conditions (p > 0.05, **Supplemental Figure 2B**). Error bars represent standard error and statistical tests are Student’s t-tests, with significance denoted by not significant (ns) p > 0.05, * p < 0.05, ** p < 0.01, and *** p < 0.001. Scale bar is 10 μm.

We also investigated whether transcription inhibition decreases nuclear blebbing through changes in chromatin-based cell nuclear rigidity, which could alter the mechanical force balance between the nucleus and cytoskeleton (Stephens *et al*., 2019b). Nuclear height measurements provide a proxy for this force balance between the nucleus and actin cytoskeleton compressing it. Using spinning disk confocal microscopy of Hoechst-stained nuclei, we measured nuclear height in untreated and VPA-treated cells without or with alpha-amanitin (**Figure 3, C and D**). VPA treatment results in a statistically significant decrease in the average height of the nucleus (**Figure 3E**). This is likely due to nuclear softening from VPA-induced chromatin decompaction (Stephens *et al*., 2018), allowing further nuclear compression by actin fibers on top of the nucleus (Khatau *et al*., 2009). Treatment with alpha-amanitin did not induce additional significant changes in nuclear height in either untreated or VPA-treated cells (**Figure 3E**) or actin contraction levels (**Supplemental Figure 1**). Thus, transcription inhibition appears to have little effect on the overall mechanical force balance between the nucleus and cytoskeleton.

To directly measure whether nuclear rigidity is altered by RNA pol II inhibition, we used micropipette micromanipulation of individual nuclei to measure the nuclear force-extension relation (Stephens *et al*., 2017; Currey *et al*., 2022). Micromanipulation force measurements uniquely provide the ability to measure both short-extension chromatin-based nuclear stiffness and lamin-A-based strain stiffening at longer extensions (>3 μm). For ease of isolation, we used MEF vimentin null (V-/-) nuclei which display similar nuclear rigidity as wild type MEFs (Stephens *et al*., 2017) and display similar nuclear blebbing behaviors (**Supplemental Figure 2A**). VPA-treated nuclei exhibited a weaker short-extension chromatin-based spring constant than untreated cell nuclei (**Figure 3, F and G**), consistent with previous reports (Stephens *et al*., 2017, 2018, 2019a). Interestingly, transcription inhibition did not alter wild type or VPA-treated chromatin-based nuclear stiffness (**Figure 3G**). This data agrees with the measurements of nuclear height inside live cells, which also did not change upon transcription inhibition **(Figure 3 E**). Nuclear strain stiffening in the long-extension lamin-dominated regime also did not significantly change across all conditions (**Supplemental Figure 2B**). However, in all transcription inhibition experiments, we observe wrinkles in the nuclear lamina (**Supplemental Figure 2C**), which can arise due to altered tension and buckling in the lamina. Altogether, we find that transcription does not increase chromatin-based nuclear rigidity and does not change overall nuclear stiffness.

### Transcription inhibition changes the type and frequency of rupture, but not the number of nuclei that rupture

Abnormal nuclear shape often correlates with the loss of nuclear compartmentalization. To determine whether decreased nuclear blebbing upon transcription inhibition also impacts nuclear rupture, we tracked nuclear ruptures in transcription-inhibited cells. Nuclear rupture can be observed by the spilling of concentrated nuclear localization signal labeled with green fluorescence protein (NLS-GFP) from the nucleus into the cytoplasm (**Figure 4A**). In live-cell imaging of NLS-GFP cells over three hours at two-minute intervals we observed that the percentage of nuclei that rupture does not change in a statistically significant manner upon transcription inhibition (**Figure 4B**). Thus, although transcription inhibition in VPA-treated cells decreased nuclear blebbing to wild type levels, it did not induce a corresponding suppression in the percentage of nuclei that undergo nuclear rupture.

**Figure 4.**
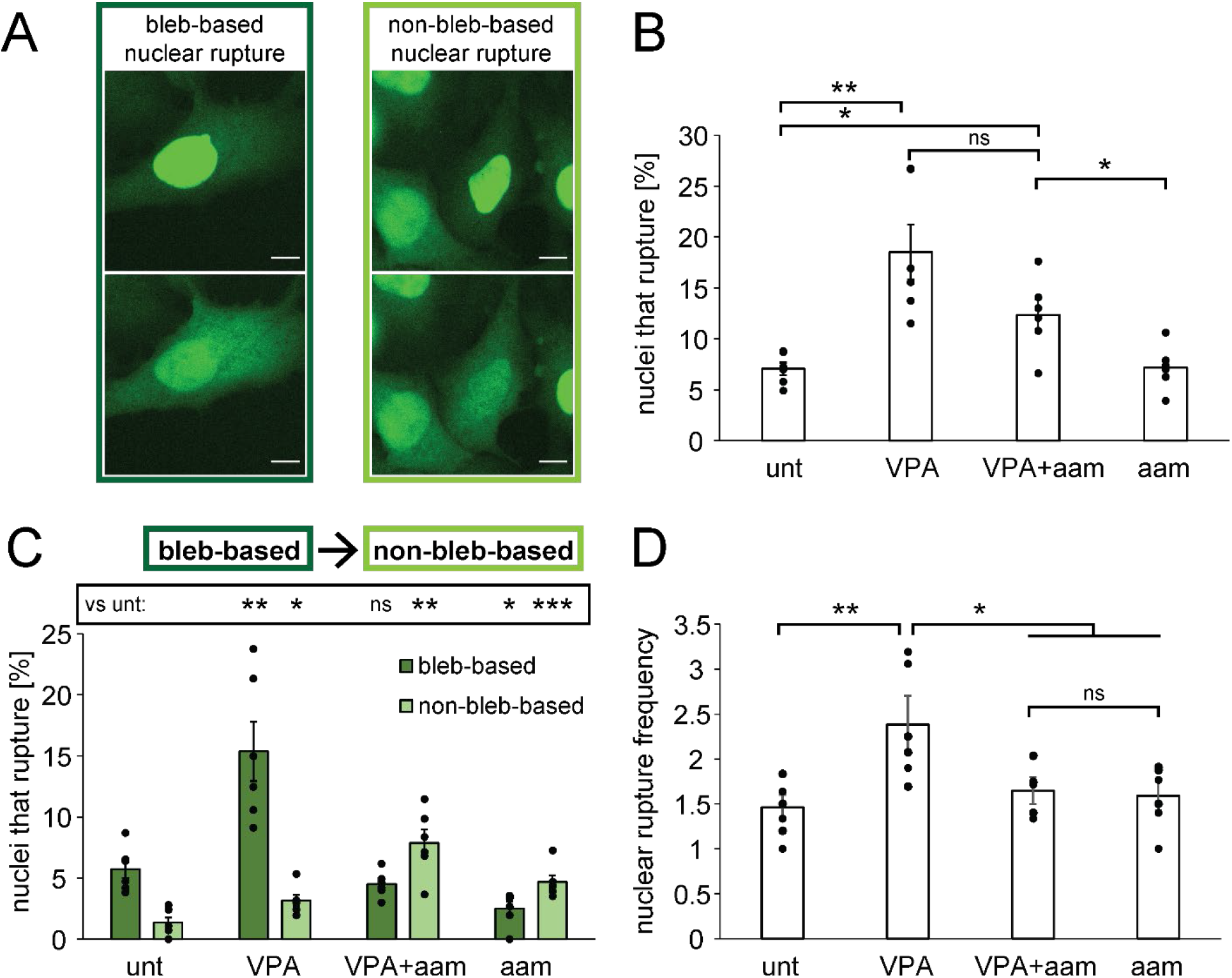
Transcription inhibition does not alter the percentage of nuclei that rupture but does alter the type and frequency of ruptures per nucleus. (A) Example images of bleb-based and non-bleb-based nuclear ruptures. (B) Graph of the percentages of nuclei that rupture in a 3-hour time-lapse with 2-minute intervals of NLS-GFP in untreated (unt), VPA-treated (VPA), VPA- and alpha-amanitin-treated (VPA+aam), and alpha-amanitin-treated cells (aam). (C) Graph of the percentages of nuclei that present bleb-based (dark green) or non-bleb-based (light green) nuclear ruptures. (D) Graph of average nuclear rupture frequency which is the number of rupture events per nucleus that displays at least one nuclear rupture. The averages were calculated from six biological replicates, graphed as dots, and each consists of n = 100-300 cells. Error bars represent standard error and statistical tests are Student’s t-tests, with significance denoted by not significant (ns) p > 0.05, * p < 0.05, ** p < 0.01, and *** p < 0.001. Scale bar is 10 μm.

Nuclear blebs have been reported to be responsible for most nuclear ruptures (Denais *et al*., 2016; Raab *et al*., 2016; Patteson *et al*., 2019; Stephens *et al*., 2019a) but ruptures can occur in non-blebbed nuclei (Robijns *et al*., 2016; Chen *et al*., 2018; Xia *et al*., 2018; Zhang *et al*., 2019). It is possible that nuclear rupture in transcription-inhibited cells occurs in non-blebbed nuclei. We therefore counted ruptures as bleb-based or non-bleb-based for untreated wild type (unt) and VPA-treated (VPA) cells without or with alpha-amanitin (aam) (**Figure 4, A and C**). In untreated wild type and VPA-treated cells with normal transcription, the majority of nuclear ruptures are bleb-based (> 80% of ruptures, dark green, p < 0.001, **Figure 4C**), in agreement with previous reports (Stephens *et al*., 2019a). Upon transcription inhibition with alpha-amanitin, the predominant type of rupture changes to non-bleb-based nuclear rupture (> 63% of ruptures, light green, p < 0.02, **Figure 4C**). A limitation of these measurements is that if a bleb formed, ruptured, and reabsorbed in less than two minutes, it would appear as a non-bleb-based rupture. However, based on the long typical lifetime of a nuclear bleb, this error is likely minimal. Thus, even though transcription inhibition suppresses nuclear blebbing, it results in a similar percentage of nuclei rupturing, the majority of which are not blebbed.

Of the nuclei that ruptured, we hypothesized that transcription might control how frequently they rupture. For each nucleus that ruptured, we counted the number of times it ruptured over the three-hour timelapse. Wild type nuclei that ruptured did so an average of 1.5 ± 0.2 times, while VPA-treated nuclei ruptured 2.4 ± 0.4 times per three hours. Dual treatment with VPA and alpha-amanitin decreased nuclear rupture frequency to wild type levels (1.6 ± 0.2 per rupturing nucleus per three hours), which was also similar to treatment with alpha-amanitin alone **(Figure 4D**). Taken together, the data shows that transcription inhibition decreases the number of times an individual nucleus may rupture, possibly by inhibiting bleb formation.

### Nuclear bleb formation and stabilization are dependent on transcription

Transcription inhibition could inhibit nuclear blebs by preventing either the formation or the stabilization of nuclear blebs. To assay these possibilities, we tracked cells during the timelapse for new bleb formation and whether that bleb stabilized or reabsorbed. Blebs were counted as stabilized if they remained after NLS-GFP reaccumulated in the nucleus, following bleb formation and rupture (**Figure 5A**, blue); alternatively, blebs were counted as reabsorbed if they disappeared following nuclear rupture (**Figure 5A,** orange). Wild type cells exhibited formation of nuclear blebs coupled to nuclear ruptures, of which the majority of the nuclei stabilized to remain blebbed, while a minority were quickly reabsorbed, with the nucleus returning to a normal shape (**Figure 5**). VPA-treated cells exhibited similar behavior, but with more nuclear bleb formation than in wild type cells (**Figure 5B**). Thus, in cells with normal transcription (unt and VPA), regardless of the level of blebbing, the majority of nuclear blebs that formed were stabilized after formation (**Figure 5C,** blue). However, transcription inhibition by alpha-amanitin drastically decreased nuclear bleb formation (**Figure 5B**). When a nuclear bleb did form, it was usually reabsorbed quickly, before NLS-GFP that spilled into the cytoplasm reaccumulated in the nucleus (**Figure 5C,** orange). Thus, both nuclear bleb formation and stabilization decreased drastically upon transcription inhibition, suggesting that transcription is a key contributor to nuclear blebs.

**Figure 5.**
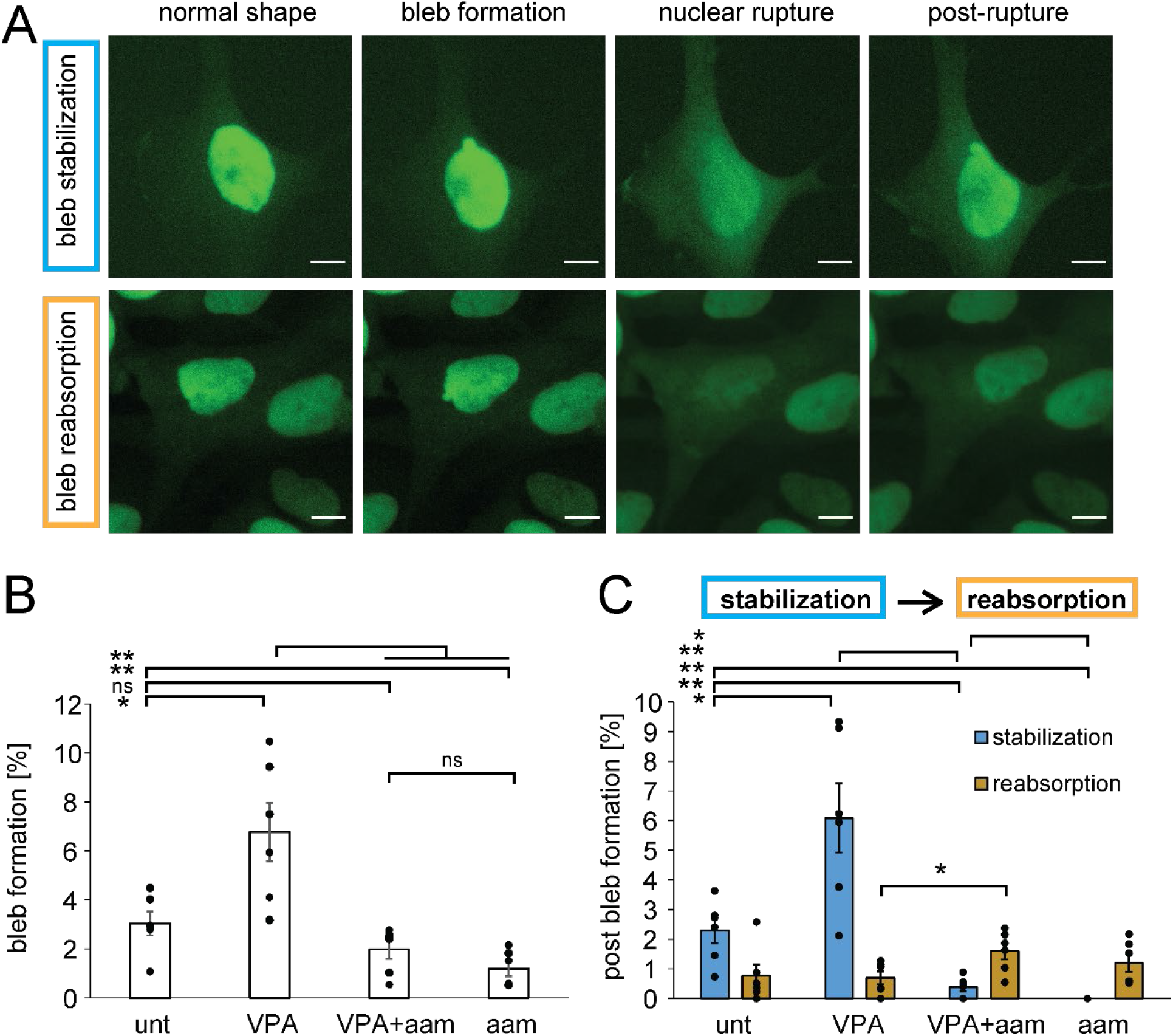
Bleb formation and stabilization are dependent on transcription. (A) Example images of new bleb formation stabilization (top, blue) or reabsorption (bottom, orange) tracked via NLS-GFP live cell imaging. (B) Graph of the percentages of nuclei that display new nuclear bleb formation in a 3-hour time-lapse for untreated (unt), VPA-treated (VPA), VPA- and alpha-amanitin-treated (VPA+aam), and alpha-amanitin-treated cells (aam). (C) Graph splitting the percentages of nuclei that display new nuclear bleb formation alongside of a nuclear rupture that results in either bleb stabilization (blue) or reabsorption (orange) post rupture. The averages were calculated from six biological replicates, graphed as dots, and each consists of n = 100-300 cells. Error bars represent standard error and statistical tests are Student’s t-tests, with significance denoted by not significant (ns) p > 0.05, * p < 0.05, ** p < 0.01, and *** p < 0.001. Scale bar is 10 μm.

### Transcriptional contribution to nuclear bleb composition

To assess how transcription aids formation and stabilization of nuclear blebs, we assayed for markers of transcription in the bleb. Previous reports indicate that active/phosphorylated RNA pol II is enriched in nuclear blebs (Shimi *et al*., 2008; Helfand *et al*., 2012; Bercht Pfleghaar *et al*.,2015). Using immunofluorescence, we assayed the distribution of active RNA pol II γSer5 (initiation) and γSer2 (elongation) relative to bulk chromatin, labeled by Hoechst (**Figure 6, A and B**). Specifically, we measured the average intensity of each transcription marker and chromatin in the nuclear bleb relative to the nuclear body to compute a ratio. Hoechst intensities reveal that on average there is less chromatin in the bleb relative to the body across all conditions (~66%, **Figure 6, A and B**), in agreement with previous reports (Bercht Pfleghaar *et al*., 2015; Stephens *et al*., 2018). Interestingly, RNA pol II γSer5 marking transcription initiation is significantly enriched in the bleb as compared to the nuclear body relative to Hoechst in all conditions (**Figure 6, A and B**). On the other hand, RNA pol II γSer2 marking transcription elongation has a decreased intensity in the bleb as compared to the nuclear body, similar to the corresponding ratio for Hoechst. Thus, transcription initiation is enriched in the bleb relative to chromatin and transcription elongation.

**Figure 6.**
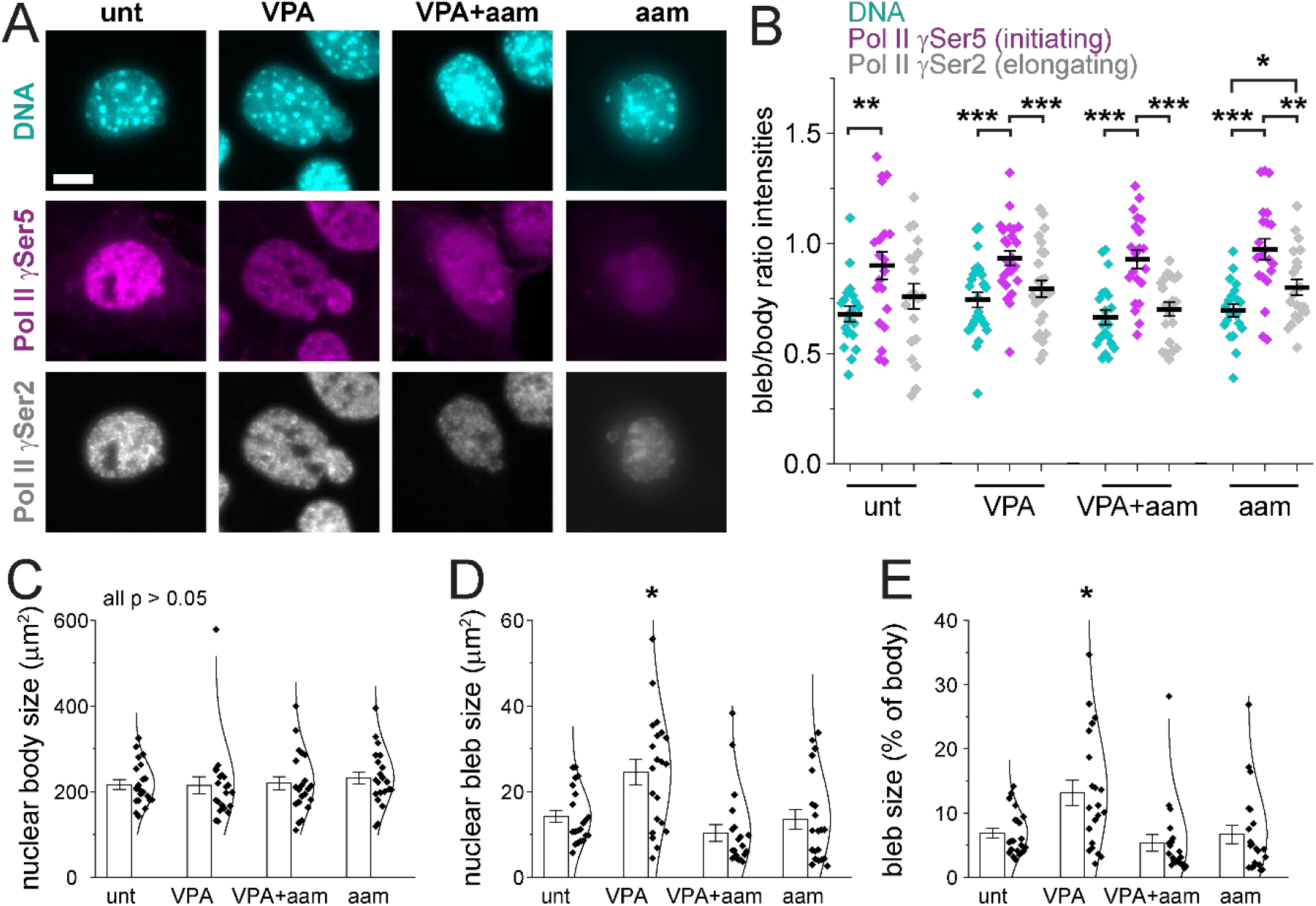
Transcription initiation is enriched in blebs relative to DNA and overall supports bleb size. (A) Example images of Hoechst (DNA, cyan), pol II phosphorylated at Ser5 (initiating, γSer5, magenta) and pol II phosphorylated at Ser2 (elongating, γSer2, gray) in untreated (unt), VPA-treated (VPA), VPA- and alpha-amanitin-treated (VPA+aam), and alpha-amanitin-treated (aam) cells. (B) Graph of the single nucleus bleb/body ratio of immunofluorescence signal. DNA (Hoechst) is decreased in the bleb, but pol II γSer5 in the bleb is enriched relative to DNA and has a similar signal as the main nuclear body. Graphs of average (C) nuclear body size, (D) nuclear bleb size, and (E) average size of bleb as a percentage of the nuclear body. (unt, n = 20; VPA, n = 26; VPA+aam, n = 21; aam, n = 21). Error bars represent standard error and statistical tests are Student’s t-tests, with significance denoted by * p < 0.05, ** p < 0.01, and *** p < 0.001. Scale bar is 10 μm.

To determine if transcription supports nuclear blebs, we assayed nuclear bleb size. The area of the nuclear body and bleb were measured in all conditions. The average size of the nuclear body did not change between untreated wild type and VPA-treated cells without or with alpha-amanitin (**Figure 6C**). However, from wild type (low nuclear blebbing) to VPA-treated cells (high nuclear blebbing), there was a significant increase in the size of nuclear blebs, suggesting that chromatin decompaction by histone acetylation can grow and stabilize blebs (**Figure 6D**). Transcription inhibition by alpha-amanitin suppresses bleb size in VPA-treated cells, returning the average size to untreated wild type levels (**Figure 6D**). Analysis of nuclear bleb size as a percentage of the body size reveals the same trends, further supporting that changes in bleb size are not due to differences in nuclear body size (**Figure 6E**). Altogether, this data suggests that transcription is essential for nuclear bleb growth and maintenance of size upon chromatin decompaction via VPA.

### Transcriptional motor activity generates nuclear deformations in active polymer simulations

To further understand the mechanism underlying transcription-mediated nuclear deformations, we developed a Brownian dynamics polymer simulation model of the nucleus. Building on previous mechanical and morphological models (Banigan *et al*., 2017; Stephens *et al*., 2017; Liu *et al*., 2021), our model is a deformable elastic, polymeric shell representing the nuclear lamina (**Figure 7A**, purple) that encapsulates a polymer chain representing the chromatin fiber (**Figure 7A**, blue). The polymer chain is crosslinked by springs to model gel-like chromatin (Strickfaden *et al*., 2020; Strom *et al*., 2021) (**Figure 7A**, red). Polymer and shell subunits are also linked by springs (**Figure 7A**, green); these linkages lead to a stiffer nucleus in simulations (Strom *et al*.,2021) and *in vivo* (Schreiner *et al*., 2015). We include RNA pol II motor activity to model transcription, which can drive motion of micron-sized chromatin domains in experiments (Zidovska *et al*., 2013; Shaban *et al*., 2018), possibly through the forces that it exerts on chromatin (Liu *et al*., 2021). In our model, RNA pol II act as extensile motors, repelling nearby chromatin, without directly interacting with the lamina **(Figure 7A**, orange, inset). Further simulation details and parameters are given in **Table 1** and the Methods section.

**Figure 7.**
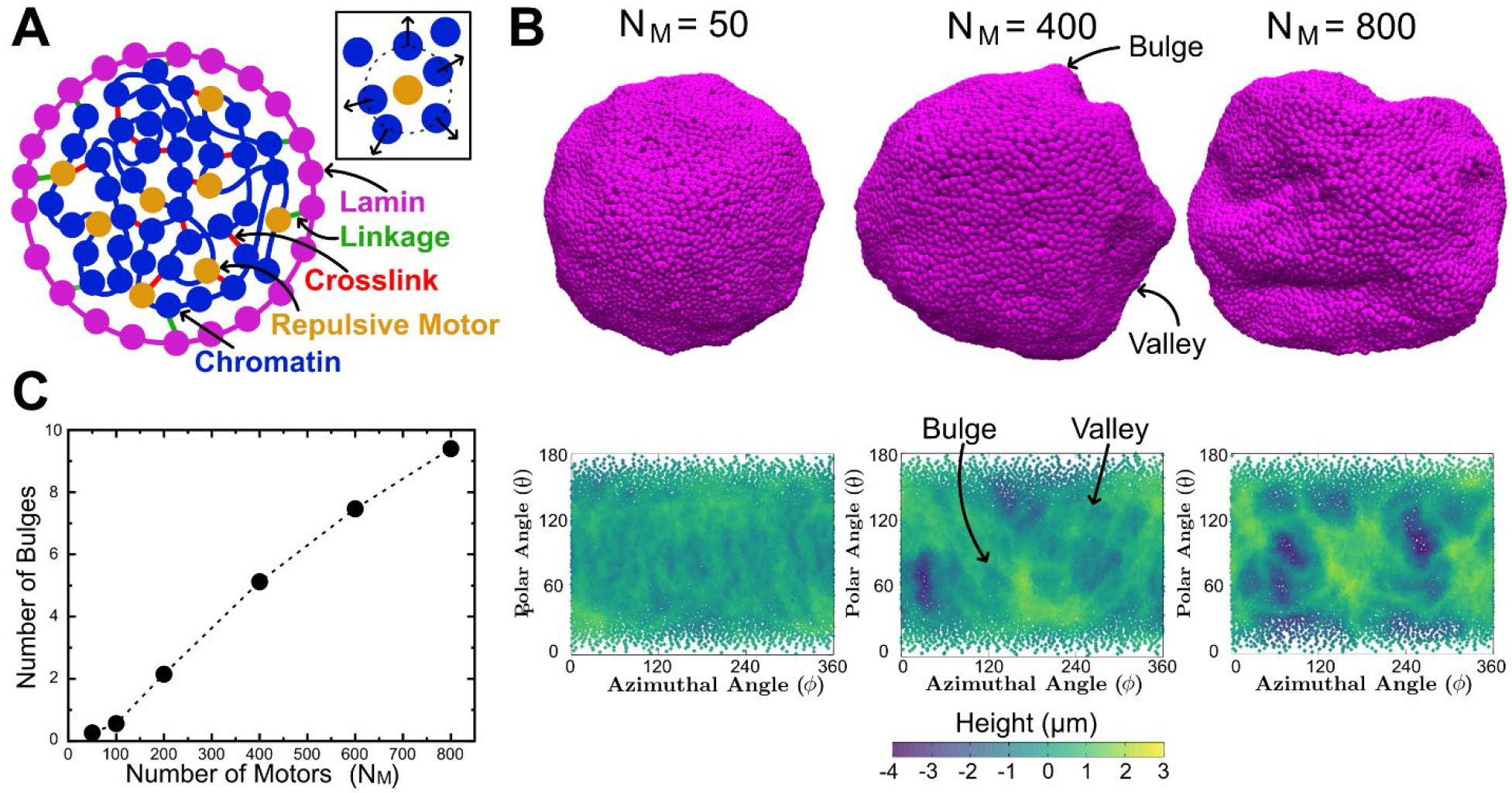
Inhibition of motors decreases nuclear bulge formation. (A) Schematic two-dimensional cross-section of the simulation model. *Inset:* Illustration of a repulsive motor (orange) repelling chromatin subunits within the interaction range. (B) *Top:* Simulation snapshots of simulated nuclei with bulges and valleys for simulations with different numbers, *N_M_*, of motors. *Bottom:* Lamina height maps corresponding to the simulation snapshots, showing bulges (green) and valleys (blue). Maps show deviation of the lamina from the mean shell radius at coordinates given by the polar angle *θ* and azimuthal angle *ϕ*. (C) Mean number of bulges increases with increasing numbers of motors.

**Table 1.** Table of simulation parameters. Parameter values used in simulations unless otherwise specified.

| Parameter | Symbol | Value |
| --- | --- | --- |
| Nuclear radius | $R$ | 10 $\mu\text{m}$ |
| Number of chromatin beads | $N$ | 5000 |
| Number of lamin beads | $N_s$ | 10000 |
| Chromatin packing fraction | $\phi$ | 0.4 |
| Number of chromatin crosslinks | $N_C$ | 2500 |
| Number of chromatin-shell links | $N_L$ | 400 |
| Number of motors | $N_M$ | 50-800 |
| Motor turnover time | $\tau$ | 10 s |
| Motor interaction range | $\sigma_m$ | 0.65 $\mu\text{m}$ |
| Chromatin spring constant | $K_{\text{chain}}$ | $1.4 \cdot 10^{-4}$ nN/$\mu\text{m}$ |
| Lamin spring constant | $K_{\text{shell}}$ | $2.8 \cdot 10^{-4}$ nN/$\mu\text{m}$ |
| Repulsion spring constant | $K_{\text{repel}}$ | $1.4 \cdot 10^{-4}$ nN/$\mu\text{m}$ |
| Diffusion constant | $D$ | 0.37 $\mu\text{m}^2/\text{s}$ |
| Bead radius | $\sigma_0$ | 0.43 $\mu\text{m}$ |
| Simulation time step | $dt$ | $5 \cdot 10^{-5}$ s |

Since alpha-amanitin inhibits RNA pol II motor activity, we performed simulations with different numbers of motors, *N_M_*. For most values of *N_M_*, simulated nuclei exhibited transient bulges, which often appeared similar to blebs and could be precursors to longer-lived blebs (**Figure 7B,** top row). To quantify this observation, we constructed height maps of the lamina (**Figure 7B**, see Methods) and identify bulges as protrusions with maximum height greater than 1 μm above the mean lamina height. Decreasing the number of motors, *N_M_*, leads to fewer bulges **(Figure 7B)** and greater homogeneity in lamina heights **(Figure 7C**). This observation indicates that chromatin motions generated by motor activity induce nuclear bulges. Inversely, inhibiting transcriptional motors suppresses bulges. These results recapitulate the experimental findings that RNA pol II activity is needed to increase nuclear blebbing (**Figure 2**), and they suggest that alpha-amanitin might suppress bleb formation by reducing the prevalence of chromatin-driven bleb precursors.

## Discussion

Chromatin is a major mechanical component of the nucleus that aids in the maintenance of both nuclear shape and compartmentalization to protect genomic organization and function. Our experiments show that the genomic process of transcription is an essential component of chromatin-driven nuclear bleb formation and bleb stabilization (**Figures 2** and **5**). However, transcription and its inhibition do not govern nuclear morphology by changing bulk nuclear rigidity (**Figure 3C-G**). Rather, transcription appears to modulate bleb formation and dynamics through an alternative mechanism, which is evident in nuclei that are already less mechanically resilient due to chromatin decompaction. Transcription initiation in particular appears to be a key component of nuclear blebs, as it is enriched in blebs (**Figure 6A-B**). Nuclear ruptures, on the other hand, are largely governed by the mechanical properties of chromatin, as transcription inhibition alters rupture frequency in individual nuclei, but not necessarily its prevalence in the entire population (**Figure 4)**. Simulations of the nucleus as a polymeric lamina with an active chromatin polymer interior suggest that blebs could arise due to the forces that transcription exerts on chromatin, which can manifest as larger chromatin motions and nuclear bulges (**Figure 7**). These findings demonstrate that transcription is a major contributor to nuclear shape, and they suggest that nuclear shape may be altered by chromatin dynamics, rather than by chromatin mechanics alone.

### Transcription is necessary for bleb formation in perturbed nuclei

Our investigation shows that transcription is necessary for excess nuclear blebbing in cells with nuclear perturbations. While nuclear blebbing increases with VPA treatment or lamin B1 knockout (LMNB1-/-), this increase is negated by transcription (**Figure 2**). Apparently, the reduction in nuclear rigidity due to either histone hyperacetylation (VPA, **Figure 3**) or histone demethylation (associated with LMNB1−/− (Stephens *et al*., 2018)) alone is insufficient to increase blebbing dramatically above wild type levels. Instead, the effects of transcription on genome spatiotemporal structure and/or dynamics are required.

Interestingly, however, transcription inhibition appears to have no effect on the basal level of nuclear blebbing present in both MEF and HT1080 cells (**Figure 2**). This data is consistent with the existence of multiple factors and processes that can drive nuclear shape disruptions, even in wild type cells. Many reports show that both actin confinement and contraction are also vital to nuclear bleb formation and stabilization (Le Berre *et al*., 2012; Hatch and Hetzer, 2016; Mistriotis *et al*., 2019). However, our data shows that neither actin confinement, as measured by nuclear height (**Figure 3E**), nor actin contraction, as measured by myosin activity (γMLC2 immunofluorescence; **Supplemental Figure 1**), are altered by transcription inhibition. Therefore, it is likely that these mechanisms can act independently, even if they also act synergistically in some scenarios.

Our findings strengthen previous reports that alterations in transcription could drive aberrant nuclear architecture in perturbed cell types. Two previous studies of cancer cell lines linked activation hormones that increase transcription to nuclear blebbing and shape abnormalities (Helfand *et al*., 2012; Chi *et al*., 2022). In prostate cancer cells (LNCaP), stimulation of transcription by testosterone-activated androgen receptor resulted in a threefold increase in nuclear blebbing, correlating with an increase in the Gleason grade of the cancer (Helfand *et al*.,2012). Similarly, in hepatocarcinoma cells (Huh7), TGFβ1 treatment resulted in the majority of nuclei becoming abnormally shaped (Chi *et al*., 2022). However, unlike in our investigation, nuclear shape changes arising after transcription induction in previous studies occurred in concert with altered expression of lamins and reorganization of chromatin and the lamina, which could affect nuclear mechanical response. Consistently, LNCaP prostate cancer cells have been reported to be softer than wild-type RWPE-1 (Khan *et al*., 2018), but required transcription upregulation to induce greater levels of nuclear blebbing. Beyond these factors, TGFβ1 induction also stimulates actin contractility (Inoue-Mochita *et al*., 2015), which is linked to nuclear deformations (Mistriotis *et al*., 2019). In contrast, we observe that histone acetylation (H3K9ac), lamin A/C, and actin contractility (γMLC2) are largely unaltered by transcriptional changes (**Figure 3A-B** and **Supplemental Figures 1–2**). Thus, our experiments and simulations suggest that even normal transcription can stimulate nuclear blebbing, and that there is an additional mechanism linking transcription to nuclear shape, beyond what has been reported previously. We propose that in parallel to previously observed pathways, transcription-induced changes to chromatin dynamics may lead to the observed nuclear shape disruptions.

### Modeling nuclear bleb formation of transcriptionally active nuclei

Our simulations of the nucleus as a polymeric lamina shell enclosing a transcriptionally active, crosslinked chromatin polymer show how transcriptional regulation of genome dynamics could impact nuclear shape. As demonstrated by our experiments, chromatin and transcription are critical components regulating nuclear shape and rupture. Models accounting only for nuclear lamins (Wren *et al*., 2012; Funkhouser *et al*., 2013) are unable to explain nuclear shape disruptions linked to chromatin compaction and/or transcription. To that end, our model provides a chromatin-based mechanism for nuclear shape disruption (**Figure 7**), while also matching other mechanical and dynamical observations of chromatin (Banigan *et al*., 2017; Stephens *et al*., 2017; Liu *et al*., 2021; Strom *et al*., 2021).

However, our model captures large nuclear deformations in the form of transient bulges rather than long-lived blebs, which leaves open the question of precisely how blebs form. While large bulges are similar to blebs, most bulges that we observe may be precursors to blebs, which could potentially form in a more complicated model with additional features. Notably, linkages, including lamin-lamin bonds, chromatin crosslinks, and chromatin-lamina links, are permanent in our model, and cannot break. *In vivo*, these linkages are dynamic and breakable under force (e.g., (Cheutin *et al*., 2003; Festenstein *et al*., 2003; Kind *et al*., 2013; Sapra *et al*., 2020; Vahabikashi *et al*., 2022)), which could contribute to the material failure that leads to nuclear bleb formation. Furthermore, we simulate isolated nuclei, which are not subject to external forces or confinement from the cytoskeleton, which can induce blebbing (Le Berre *et al*., 2012; Hatch and Hetzer, 2016; Mistriotis *et al*., 2019). Inclusion of these features could reconcile our model with experiments and allow bulges to develop into blebs.

### Nuclear mechanics dictates nuclear rupture but not necessarily shape

Surprisingly, we find that while nuclear stiffness largely controls nuclear rupture, it is not the sole determinant of nuclear shape. This contrasts with previous studies, which suggested that the nucleus maintains its shape by resisting cytoskeletal and/or other external antagonistic forces (Khatau *et al*., 2009; Le Berre *et al*., 2012; Hatch and Hetzer, 2016; Stephens *et al*., 2018; Earle *et al*., 2020). Thus, loss of chromatin- and/or lamin-based rigidity could cause the nucleus to succumb to cytoskeletal forces and form a nuclear herniation or bleb. Since transcription is needed to form and stabilize nuclear blebs, at least some aspect of nuclear shape deformations appears to be non-mechanical.

In contrast, force balance between the nucleus and the cytoskeleton appears to control nuclear rupture, even in scenarios where transcription is disrupted and blebs are not present **(Figure 4A-C).** Regardless of the transcriptional state of the nucleus, nuclei with softer, decompacted chromatin (VPA-treated) were more likely to rupture, with or without blebs. Nonetheless, the force balance governing nuclear rupture is apparently affected, but not controlled by the presence of blebs, as nuclei with blebs ruptured more frequently (*i.e*., multiple times in several hours; **Figure 4D**). Therefore, transcriptionally active nuclei that undergo rupture may suffer greater DNA damage (Stephens, 2020) or disruption of the cell cycle (Pfeifer *et al*., 2018). Furthermore, the higher frequency of ruptures agrees with reports that high nuclear curvature, such as that found on the surface of blebs, promotes rupture (Robijns *et al*., 2016; Stephens *et al*., 2018; Xia *et al*., 2018; Nmezi *et al*., 2019; Pfeifer *et al*., 2022), perhaps resulting in higher surface tensions in the lamina (Deviri *et al*., 2017; Xia *et al*., 2018; Srivastava *et al*., 2021) Therefore, although high nuclear curvature can promote rupture, it is not necessary.

However, high nuclear curvature is not necessary for rupture. This is demonstrated by our experiments showing that most nuclear ruptures in transcription-inhibited cells occur in non-blebbed and relatively normally shaped nuclei **(Figure 4C**). The high curvature of blebs was also found to be dispensable in previous studies observing ruptures without blebbing or substantial nuclear shape fluctuations (Robijns *et al*., 2016; Chen *et al*., 2018; Penfield *et al*., 2018; Earle *et al*., 2020). This notion suggests that local tensile stress can induce nuclear ruptures without necessarily leading to abnormal nuclear shape or blebbing (Zhang *et al*., 2019). We therefore argue that in VPA-treated cells with transcription inhibited, chromatin is soft such that it does not support the lamina against rupture via cytoskeletal forces, but the loss of transcription prevents the formation and stabilization of blebs.

### The mechanism of transcriptional regulation of nuclear shape

Since transcription promotes blebbing but does not alter whole-nucleus stiffness, we propose that an alternative mechanism connects these phenomena. Our experiments and modeling suggest that transcription might promote nuclear blebbing by increasing chromatin-driven pushing and pulling on the nuclear lamina.

These effects could arise through the correlated motions of micron-sized chromatin domains and/or chromatin density fluctuations that emerge due to transcriptional activity (Zidovska *et al*.,2013; Shaban *et al*., 2018; Liu *et al*., 2021). In this scenario, transcription could sporadically drive large regions of chromatin into the lamina, leading to bulges and possibly blebs. Consistently, previous simulations of isolated nuclei showed that (transcriptional) motor activity in an active, crosslinked polymer can drive correlated polymer (chromatin) dynamics and enhanced (nuclear) shape fluctuations (Liu *et al*., 2021).

Additionally, polymeric shells, such as the nuclear lamina, subject to tensile or compressive stresses can buckle, thereby undergoing a sudden, dramatic change in shape resembling a first-order phase transition (Paulose and Nelson, 2013; Yong *et al*., 2013; Banigan *et al*., 2017) Thus, there may generally be a kinetic barrier to bleb formation, which can be overcome in part by energy generated by transcriptional dynamics.

Complementarily to these dynamic mechanisms, chromatin linkages to the lamina may play a role in governing bleb formation. A previous study of lamina-associated chromatin domains (LADs) showed that transcription of genes tends to detach them from the lamina, while transcription inhibition induces lamina attachment of inactivated genes (Brueckner *et al*., 2020). Based on previous simulations suggesting that chromatin-lamina linkages can help maintain nuclear shape (Banigan *et al*., 2017; Lionetti *et al*., 2020), it is possible that linkages induced by transcription inhibition suppress bleb formation. Nonetheless, regulation of linkages might be only a secondary mechanism, since we do not observe any change in nuclear rigidity upon transcription inhibition (**Figure 3**), as seen in previous simulations (Strom *et al*., 2021).

It has also been proposed that transcription affects chromatin-chromatin connections (Nagashima *et al*., 2019), which in turn could alter nuclear rigidity, and potentially nuclear shape. Experiments imaging single-nucleosomes *in vivo* showed that transcription inhibition by alpha-amanitin or DRB enhanced spatial fluctuations of nucleosomes, suggesting that there were fewer constraints within transcriptionally inactive chromatin. Notably, inhibition by actinomyosin D had the opposite effect of suppressing nucleosome fluctuations. We find that transcription inhibition by alpha-amanitin does not reduce nuclear rigidity (**Figure 3**), as we would expect if chromatin crosslinking or bridging by RNA pol II has nucleus-scale effects (Stephens *et al*., 2017; Strom *et al*., 2021). Therefore, it is likely that transcriptional effects on crosslinking are not a major contributor to nuclear shape.

Another clue about the mechanism is the composition of the nuclear bleb. The transcription initiation marker RNA pol II γSer5 is enriched relative to DNA, while the transcription elongation marker RNA pol II γSer2 is not (**Figure 6**). This is a novel finding that clarifies which active RNA pol II is prominently enriched (Shimi *et al*., 2008; Helfand *et al*., 2012; Bercht Pfleghaar *et al*.,2015). This suggests that components associated with transcription initiation are important for bleb stabilization and growth, and possibly, formation. However, the contribution of transcription initiation is presently unclear.

Nonetheless, our results emphasize the importance of chromatin and its constituents in regulating nuclear shape, even in situations where other nuclear mechanical components, such as lamins, are unaltered. More generally, regardless of the precise physical mechanism, our study raises the possibility that other chromatin-bound molecular motors, such as condensin, cohesin, topoisomerase, or DNA polymerase, might influence nuclear shape through their activities.

## Methods

### Cell Culture

MEF WT, V-/-, LMNB1−/− and HT1080 cells were grown in DMEM (Corning) complete with 1% Pen Strep (Fisher Scientific) and 10% fetal bovine serum (FBS; HyClone) at 37°C and 5% CO2. Cell culture glass bottom 4-well imaging dishes were prepared 48 hours before imaging. Cells were passaged every other day into fresh DMEM complete. 100μl of MEF-WT and MEF-NLS-GFP cells were added into the corner of each well and 600μl of DMEM complete was added carefully in order to flow over the cells and fill the well. Drug treatment was done 22-24h prior to imaging. VPA was added to a final concentration of 2mM (20mM stock solution) and aam at 10μM (1mM stock solution). For actinomycin D (amD) treatment, amD was added at a concentration of 10μg/ml with an incubation time of no more than 30 minutes prior to imaging.

### Imaging

Images were acquired with Nikon Elements software on a Nikon Instruments Ti2-E microscope with Crest V3 Spinning Disk Confocal, Orca Fusion Gen III camera, Lumencor Aura III light engine, TMC CleanBench air table, with 40x air objective (N.A 0.75, W.D. 0.66, MRH00401) or Plan Apochromat Lambda 100x Oil Immersion Objective Lens (N.A. 1.45, W.D. 0.13mm, F.O.V. 25mm, MRD71970). Live cell time lapse imaging was possible using Nikon Perfect Focus System and Okolab heat, humidity, and CO2 stage top incubator (H301). Images were captured via camera 16 bit for population images or 12 bit sensitive for time lapse live cell imaging with 40x air objective N.A 0.75 (Nikon MRH00401). For time lapse data, images were taken in 2-minute intervals during 3 hours with 9 fields of view for each condition.

### Bleb Count

MEF cells were treated with Hoechst at a dilution of 1:20,000 to 1:40,000 for 15 minutes before population imaging or imaged via NLS-GFP to provide nuclear shape. Images were taken with 9 fields of view for each condition, more than 100 nuclei were counted, and the percentage of cells showing blebbed nuclei calculated (> 100) for each biological replicate (≥ 3). Nuclei were scored as blebbed if a protrusion 1 μm in diameter or larger was present, as previously outlined in (Stephens *et al*., 2018).

### Nuclear Rupture Analysis

MEF NLS-GFP cells were used. Image stacks were analyzed using the NIS Elements AR Analysis software. For each condition, the total number of nuclei was counted at the first and last frame of the time lapse and averaged. The number of blebbed nuclei was counted as any nuclei that displayed a nuclear bleb at any time during the time lapse. Nuclear ruptures were determined by a > 25% change in the NLS-GFP intensity cytoplasm/nucleus using 5 μm x 5 μm box in each and background subtracted. The total numbers of nuclei showing a nuclear rupture were counted differentiating between bleb-based ruptures and non-blebbed ruptures. Bleb-based ruptures were defined as nuclei showing a bleb prior to rupturing whereas non-blebbed ruptures did not show a bleb on the nucleus. The count of ruptures for each rupturing nucleus was noted for rupture frequency calculations which is the average number of ruptures per nuclei showing a nuclear rupture. Additionally, the number of blebs formed during the time lapse was counted differentiating between first: blebs forming, rupturing, and thereby stabilizing the bleb, and second: blebs forming but disappearing without showing a nuclear rupture or blebs forming, rupturing but disappearing and not being stabilized by the rupture. This analysis justifies counting the percentage of blebbed cells only at one or two timepoints as the dynamics of the formation of new blebs is captured. For each condition, 3 fields of views were analyzed for each replicate. Graphs were made in order to show the percentage of ruptures as the total number of ruptured nuclei per total number of nuclei in each field of view.

### Nuclear Height Analysis

The nuclei were imaged using a Spinning Disk Confocal with a Plan Apochromat Lambda 100x Oil Immersion Objective Lens (N.A. 1.45, W.D. 0.13mm, F.O.V. 25mm). 77 images with an axial distance of 0.2 μm were taken as z-slices with a total distance of 15μm to be covered. The center of one nucleus at a time was selected and a slices view was created. The fluorescence intensity profile for the z-slice was analyzed by using a full-width half-maximum (FWHM) calculation giving the width of the fluorescence intensity peak corresponding to the height of the nucleus. Two measurements of nuclear height were averaged for each nucleus. 10-20 nuclei were measured for each condition.

### Micromanipulation Force Measurements

As first described (Stephens *et al*., 2017) and more recently updated (Currey *et al*., 2022), MEF vimentin null (V-/-) cells were grown in a micromanipulation well to provide low angle access via micropipettes. MEF V−/− nuclei were isolated from living cells via spray micropipette of mild detergent Triton X-100 0.05% in PBS. The pull micropipette was used to grab the nucleus. The isolated nucleus was then grabbed at the opposite end with a precalibrated force micropipette and suspended in preparation for force-extension measurements. The pull pipette was moved at 50 nm/sec to provide 3 or 6 μm extension to the nucleus. The pull micropipette is tracked to provide nucleus extension (μm) while measurement of the deflection of the force micropipette multiplied by the bending modulus (1.2 - 2 nN/μm) provides the measure of force (nN). The slope of the force vs. extension plot provides the spring constant (nN/μm) for the short chromatin-dominated regime (< 3 μm) and long extension lamin A dominated strain stiffening regime (> 3 μm). The long regime spring constant minus the short regime spring constant provides the measure of lamin A-based strain stiffening.

### Immunofluorescence

Cells were grown in 8-well dishes for 48 hours prior to fixation. Treatment with drugs was done 24 hours prior to fixation which was done with a solution of 3.3% PFA in Trition X-100 for 15 minutes. 3 washing steps were performed with phosphate buffered saline (PBS) and the last one with PBS-Tween 20 (PBS-T). Primary antibodies were diluted in 10% goat serum in PBS (GPBS) and incubated with the cells for 1h at 37°C before washing 3 times with PBS. Secondary antibodies were incubated in GPBS for 1h at room temperature. The primary antibody used were LaminA/C at 1:200 (CST 4C11, Mouse mAb #4777), H3K9ac at 1:500 (CST C5B11 Rabbit mAb #9649), Anti-RNA polymerase II CTD repeat YSPTSPS (phospho S5) antibody at 1:1000 (4H8 - ChIP Grade, ab5408 Mouse mAb), and Anti-RNA polymerase II CTD repeat YSPTSPS (phospho S2) antibody at 1:1000 (ab5095, Rabbit pAb). Secondary antibodies were used at a 1:1000 dilution and include Alexa Fluor 488, 555, 647 Goat anti-Mouse or Goat anti-Rabbit IgG (H+L), F(ab’)2 Fragment (CST 4408-4414). The nuclei were stained using a Hoechst solution of Hoechst in PBS at a 1:40 000 dilution for at least 5 minutes. Cells were kept and imaged in PBS or mounted using Prolong Gold anti-fade mountant (Invitrogen, P36930) and let sit overnight out of light. Fluorescence intensities were analyzed by measuring single nuclei as ROI and subtracting background via a 30X30 pixel area with no cells. All single nuclei measures were averaged over multiple fields of view to provide a single average intensity measurement for each experiment, for which there were at least three replicates. For comparison, all intensities were normalized with the mean value for untreated nuclei.

For bleb vs body measurements an ROI was hand drawn around each and intensities were background subtracted as detailed above.

### RNA labeling

Labeling RNA was accomplished using Click-iT™ RNA Alexa Fluor™ 594 Imaging Kit (Invitrogen, C10330) previously described (Jao and Salic, 2008). Cells were plated in 8 well coverslips (Cellvis, C8-1.5H-N) grown and left untreated or treated with VPA and/or alpha amanitin. 1 mM final concentration of EU was added to cells for 1 hour incubation. Cells were fixed and permeabilized same as in immunofluorescence. After 2X PBS washes the 500 μL formulation of the Click-IT reaction cocktail was added and allowed to incubate for 30 minutes out of light. The reaction was terminated by removing and washing with the defined Click-IT reaction rinse buffer. The nuclei were stained using a Hoechst solution of Hoechst in PBS at a 1:40 000 dilution for at least 5 minutes. The cells were rinsed 2X more times in PBS. Cells were kept and imaged in PBS or mounted using Prolong Gold anti-fade mountant (Invitrogen, P36930) and let sit overnight out of light.

### Simulation model

For the simulation of the nuclear lamina shell, we first generated 10,000 point particles (or “subunits”) in the Fibonacci sequence on a sphere of *R* = 10 μm. Since the nuclear lamina is not a regular lattice (Shimi *et al*., 2015; Mahamid *et al*., 2016; Turgay *et al*., 2017), we then randomized their positions on the shell with thermal noise. While randomizing subunit positions, we gradually and sequentially increased the size of subunits to *σ*_0_ such that no large forces were generated due to overlap. Subunits cannot overlap due to soft repulsive potential given by:

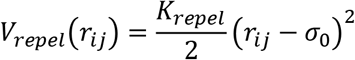

for *r_ij_*≤*σ_0_*, where *r_ij_* is the distance between particles *i* and *j*, and *K*_soft_ is the spring constant. Once the lamin subunits reach size *σ*_0_, we randomize again with the current size to ensure that no remnant of the original lattice survives in the simulated structure. *In vivo*, the nuclear lamin network has an average coordination number of approximately 4 to 5 (Shimi *et al*., 2015; Sapra *et al*., 2020). To model this, we developed an algorithm to connect the lamin subunits such that the average coordination number of the lamin network is around 4.5.

To model the chromatin chain, we first generated a random walk on a face-centered cubic lattice with no overlapping steps. We set a confining spherical boundary condition at radius *R* on the chain and subsequently decreased the chain size such that it was compacted into the sphere. The Rouse chain subunits are joined by harmonic springs, which also disfavors overlap by a soft repulsive potential:

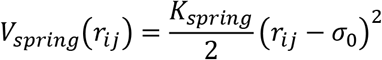

We then equilibrated the compacted chain inside the lamina shell. We modeled chromatin-chromatin crosslinks and chromatin-lamina linkages by randomly choosing nearby subunits on the chromatin chain and nuclear lamina and connecting them by harmonic springs. Purely repulsive (“extensile”) motors are simulated as generating a monopolar (outward) repulsive force on chromatin subunits if they are within the interaction range, *σ_m_*, of the motor. Chromatin motors do not exert any force on the nuclear lamin subunits.

### Simulation methods

The time evolution of the system is governed by overdamped Langevin dynamics:

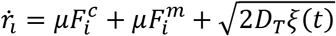

where *r_i_* is position of *i*^th^ particle; 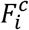 is the conservative force on *i*^th^ particle, calculated as 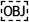; and 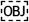 is the non-conservative force due to the repulsive motors that repel chromatin subunits radially outward. Simulation parameters are given in **Table 1**. We integrate the equation of motion by the Euler-Murayama method.

In our model, the chromatin chain has 5000 subunits and the nuclear lamina (shell) is has 10,000 subunits. There are 400 chromatin-shell linkages and 2500 crosslinkers. Each chromatin or lamin subunit has a diameter of σ=0.43089 μm. The spring constant (both *K_spring_* and *K_repel_*) for the chain is 1.4*10^-4^ nN/μm and the spring constant for the shell is 2.8*10^-4^ nN/μm. We evolved the simulation for 10^7^ timesteps, which we take to be 500 s. We ran 10 realizations for each motor number, *N_m_*, shown in **Figure 7C**, identified and counted bulges for each realization (see below), and averaged over realizations to compute the mean bulge number. Motors stochastically switch off of one subunit and onto another, random selected chromatin subunit, with a mean turnover time of 10 s, as in previous simulations (Liu *et al*., 2021).

### Identification of bulges in simulations

To identify bulges and valleys, we first calculate the average shell radius and compute the height of each shell subunit above or below this radius. Subunits with height greater than 1 μm are considered to be part of a bulge.

To identify and count bulges, we project the shell, using spherical polar coordinates, onto a 2D map of the polar and azimuthal angles of each subunit (*θ* and *ϕ*, respectively). Bulges are first identified by local maxima in subunit heights. To determine the area that a bulge occupies (and thus identify unique bulges), first centered the map on the local maximum, and then considered heights of subunits within a circle around that point. Subunits with height greater than 1 μm were counted as part of the bulge. If there were subunits within the circle that were part of the bulge, we expanded the circle radius to look for additional subunits in the bulge. After expanding the circle, if no additional nearby subunits had height greater than 1 μm, we defined the bulge as containing only the subunits that had already been counted. Bulges were defined such that each bulge contains a unique set of subunits (*i.e*., overlapping bulges were merged to be considered a single bulge).

## Acknowledgements

We would like to thank Paul Janmey for helpful discussions and Kuang Liu for helpful discussions and advice on implementing the simulation model. IKB, MLC, YB, MP, and ADS are supported by the Pathway to Independence Award (R00GM123195). AEP is supported by NIH R35GM142963. JMS is supported by NSF-DMR-1832002 and SG is funded by a Syracuse University dissertation fellowship. IKB, MLC, EJB, and ADS are supported by Center for 3D Structure and Physics of the Genome 4DN2 grant (1UM1HG011536).

---

**Supplemental Figure 1.**
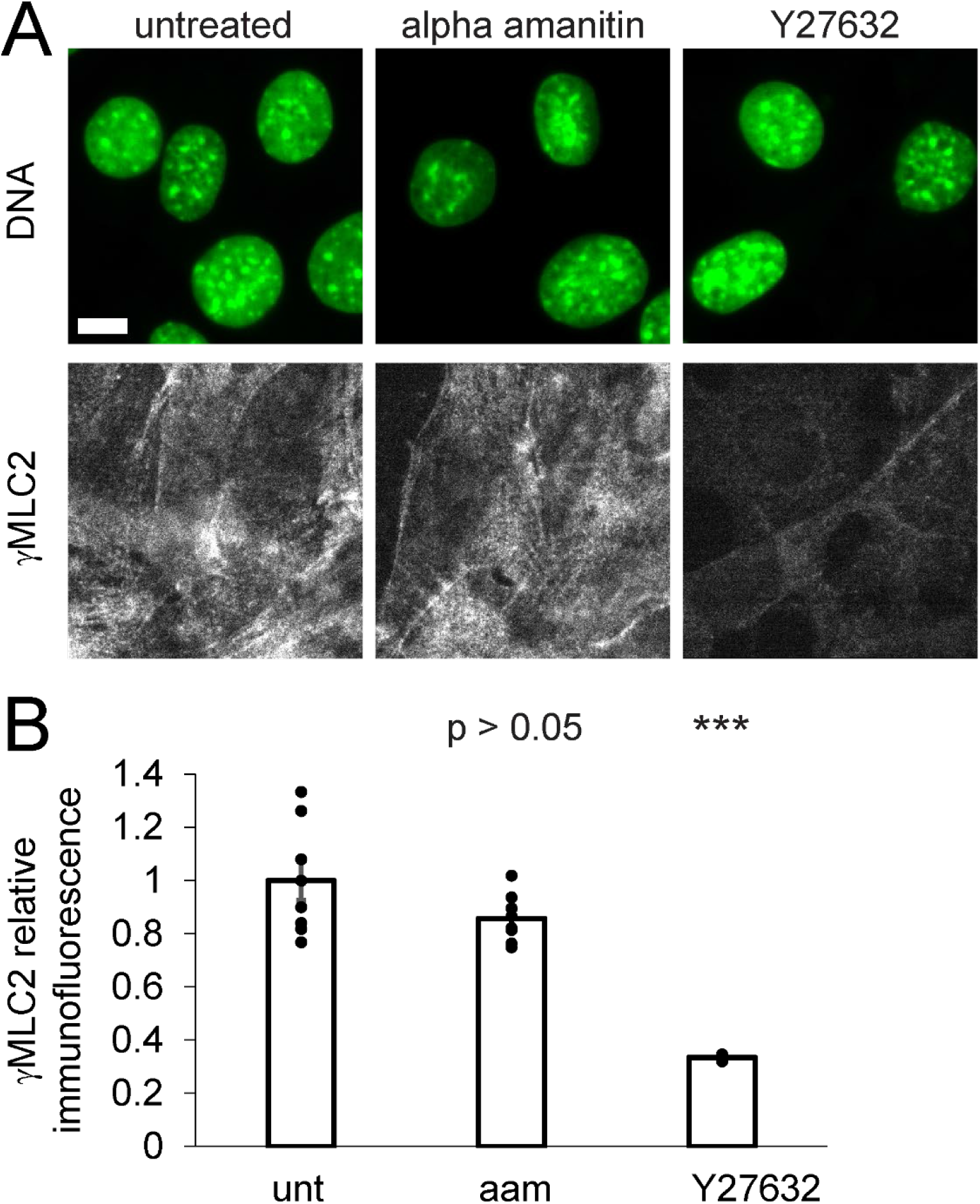
Transcription inhibition does not alter active actomyosin contraction. (A) Example immunofluorescence images of DNA via Hoechst (green) and active myosin via phosphorylated myosin light chain 2 (γMLC2, gray) for untreated (unt), alpha-amanitin (aam), and actin contraction inhibition via ROCK inhibitor Y27632. (B) Relative γMLC2 immunofluorescence for each condition. Four to eight biological replicates for untreated (unt) and alpha-amanitin (aam) populations and four biological replicates for Y27632, consisting of 30 cells each, are shown as dots. Error bars represent standard error and statistical tests are Student’s t-tests, with significance denoted by * p < 0.05, ** p < 0.01, and *** p < 0.001. Scale bar is 10 μm.

**Supplemental Figure 2.**
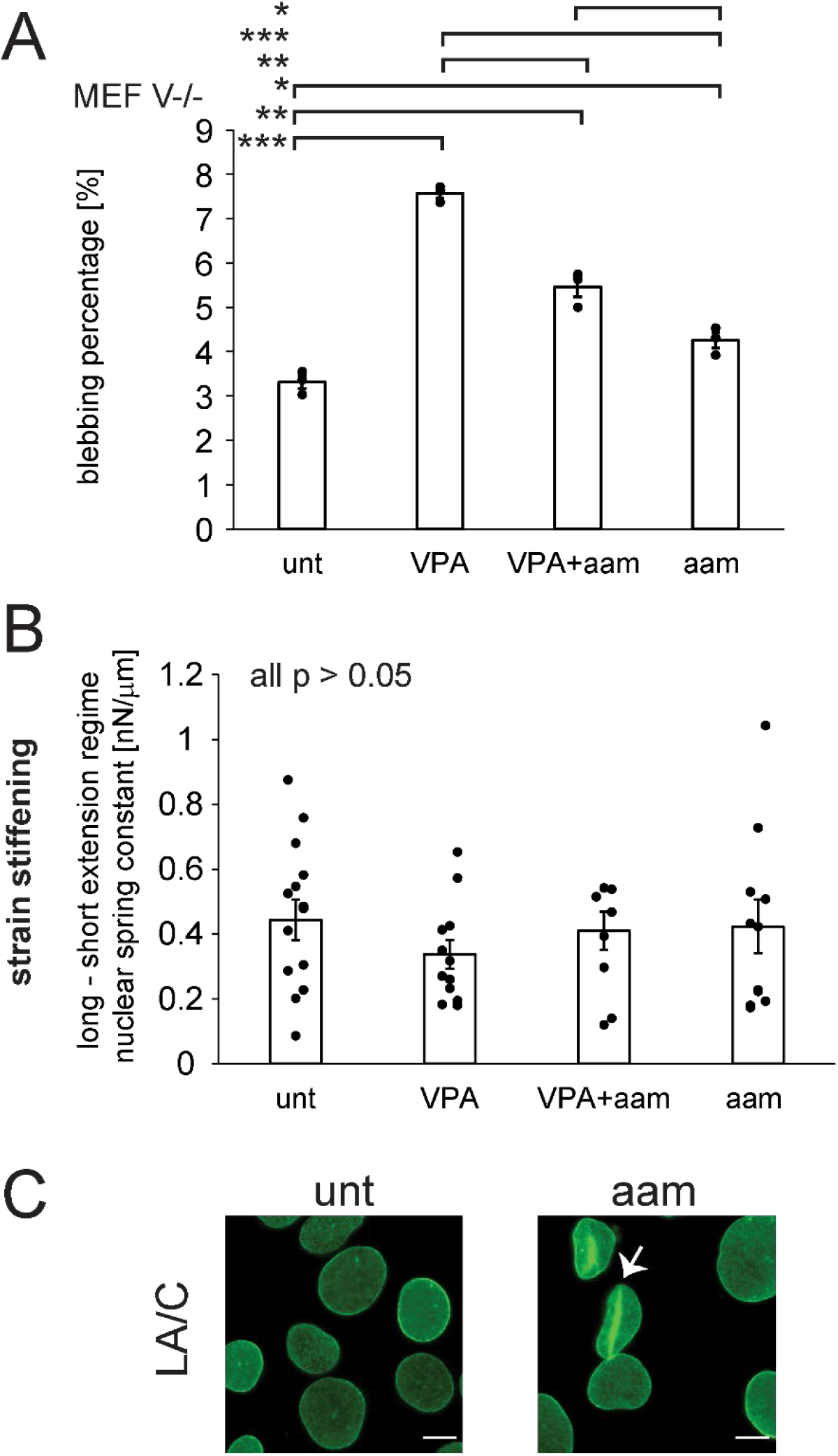
Transcription inhibition does not alter the lamin-dominated regime. (A) Graph of nuclear blebbing percentages for untreated wild-type (unt) and VPA-treated cells without or treated with alpha-amanitin (aam). Three biological replicates represented as dots consisted of > 200 cells each. (C) Graph of the lamin-A-based strain-stiffening nuclear spring constant (long regime minus short regime spring constant). unt, n = 12; VPA, n = 12; VPA+aam, n = 11; aam, n = 8. Error bars represent standard error and statistical tests are Student’s t-tests, with significance denoted by * p < 0.05, ** p < 0.01, and *** p < 0.001. (C) Example images of lamin A/C immunofluorescence showing lamin wrinkles upon alpha-amanitin treatment, denoted by white arrow. Scale bar is 10 μm.

***Supplemental Table 1. Raw Data*.** This is an excel document with compiled data used for making each figure.

## References

Banigan, EJ, Stephens, AD, and Marko, JF (2017). Mechanics and buckling of biopolymeric shells and cell nuclei. Biophys J 113, 1654–1663.

Banigan, EJ, Tang, W, van den Berg, AA, Stocsits, RR, Wutz, G, Brandão, HB, Busslinger, GA, Peters, J-M, and Mirny, LA (2022). Transcription shapes 3D chromatin organization by interacting with loop extrusion.

Belaghzal, H, Borrman, T, Stephens, AD, Lafontaine, DL, Venev, SV, Weng, Z, Marko, JF, and Dekker, J (2021). Liquid chromatin Hi-C characterizes compartment-dependent chromatin interaction dynamics. Nat Genet 53, 367–378.

Bercht Pfleghaar, K, Taimen, P, Butin-Israeli, V, Shimi, T, Langer-Freitag, S, Markaki, Y, Goldman, AE, Wehnert, M, and Goldman, RD (2015). Gene-rich chromosomal regions are preferentially localized in the lamin B deficient nuclear blebs of atypical progeria cells. Nucleus 6, 66–76.

Brueckner, L, Zhao, PA, van Schaik, T, Leemans, C, Sima, J, Peric-Hupkes, D, Gilbert, DM, and van Steensel, B (2020). Local rewiring of genome-nuclear lamina interactions by transcription. EMBO J 39, e103159.

Buenrostro, JD, Giresi, PG, Zaba, LC, Chang, HY, and Greenleaf, WJ (2013). Transposition of native chromatin for fast and sensitive epigenomic profiling of open chromatin, DNA-binding proteins and nucleosome position. Nat Methods 10, 1213–1218.

Chalut, KJ, Höpfler, M, Lautenschläger, F, Boyde, L, Chan, CJ, Ekpenyong, A, Martinez-Arias, A, and Guck, J (2012). Chromatin decondensation and nuclear softening accompany Nanog downregulation in embryonic stem cells. Biophys J 103, 2060–2070.

Chen, NY, Kim, P, Weston, TA, Edillo, L, Tu, Y, Fong, LG, and Young, SG (2018). Fibroblasts lacking nuclear lamins do not have nuclear blebs or protrusions but nevertheless have frequent nuclear membrane ruptures. Proc Natl Acad Sci U S A 115, 10100–10105.

Cheutin, T, McNairn, AJ, Jenuwein, T, Gilbert, DM, Singh, PB, and Misteli, T (2003). Maintenance of stable heterochromatin domains by dynamic HP1 binding. Science 299, 721–725.

Chi, Y-H, Wang, W-P, Hung, M-C, Liou, G-G, Wang, J-Y, and Chao, P-HG (2022). Deformation of the nucleus by TGFβ1 via the remodeling of nuclear envelope and histone isoforms. Epigenetics Chromatin 15, 1.

Chu, F-Y, Haley, SC, and Zidovska, A (2017). On the origin of shape fluctuations of the cell nucleus. Proc Natl Acad Sci U S A 114, 10338–10343.

Currey, ML, Kandula, V, Biggs, R, Marko, JF, and Stephens, AD (2022). A versatile micromanipulation apparatus for biophysical assays of the cell nucleus. Cell Mol Bioeng.

De Vos, WH et al. (2011). Repetitive disruptions of the nuclear envelope invoke temporary loss of cellular compartmentalization in laminopathies. Hum Mol Genet 20, 4175–4186.

Denais, CM, Gilbert, RM, Isermann, P, McGregor, AL, te Lindert, M, Weigelin, B, Davidson, PM, Friedl, P, Wolf, K, and Lammerding, J (2016). Nuclear envelope rupture and repair during cancer cell migration. Science 352, 353–358.

Deviri, D, Discher, DE, and Safran, SA (2017). Rupture dynamics and chromatin herniation in deformed nuclei. Biophys J 113, 1060–1071.

Earle, AJ, Kirby, TJ, Fedorchak, GR, Isermann, P, Patel, J, Iruvanti, S, Moore, SA, Bonne, G, Wallrath, LL, and Lammerding, J (2020). Mutant lamins cause nuclear envelope rupture and DNA damage in skeletal muscle cells. Nat Mater 19, 464–473.

Festenstein, R, Pagakis, SN, Hiragami, K, Lyon, D, Verreault, A, Sekkali, B, and Kioussis, D (2003). Modulation of heterochromatin protein 1 dynamics in primary Mammalian cells. Science 299, 719–721.

Funkhouser, CM, Sknepnek, R, Shimi, T, Goldman, AE, Goldman, RD, and Olvera de la Cruz, M (2013). Mechanical model of blebbing in nuclear lamin meshworks. Proc Natl Acad Sci U S A 110, 3248–3253.

Furusawa, T, Rochman, M, Taher, L, Dimitriadis, EK, Nagashima, K, Anderson, S, and Bustin, M (2015). Chromatin decompaction by the nucleosomal binding protein HMGN5 impairs nuclear sturdiness. Nat Commun 6, 6138.

Hatch, EM, and Hetzer, MW (2016). Nuclear envelope rupture is induced by actin-based nucleus confinement. J Cell Biol 215, 27–36.

Helfand, BT, Wang, Y, Pfleghaar, K, Shimi, T, Taimen, P, and Shumaker, DK (2012). Chromosomal regions associated with prostate cancer risk localize to lamin B-deficient microdomains and exhibit reduced gene transcription. J Pathol 226, 735–745.

Hnisz, D, Shrinivas, K, Young, RA, Chakraborty, AK, and Sharp, PA (2017). A phase separation model for transcriptional control. Cell 169, 13–23.

Hobson, CM, Kern, M, O’Brien, ET, 3rd, Stephens, AD, Falvo, MR, and Superfine, R (2020). Correlating nuclear morphology and external force with combined atomic force microscopy and light sheet imaging separates roles of chromatin and lamin A/C in nuclear mechanics. Mol Biol Cell 31, 1788–1801.

Hsieh, T-HS, Cattoglio, C, Slobodyanyuk, E, Hansen, AS, Rando, OJ, Tjian, R, and Darzacq, X (2020). Resolving the 3D landscape of transcription-linked mammalian chromatin folding. Mol Cell 78, 539–553.e8.

Inoue-Mochita, M, Inoue, T, Fujimoto, T, Kameda, T, Awai-Kasaoka, N, Ohtsu, N, Kimoto, K, and Tanihara, H (2015). p38 MAP kinase inhibitor suppresses transforming growth factor-β2-induced type 1 collagen production in trabecular meshwork cells. PLoS One 10, e0120774.

Irianto, J, Pfeifer, CR, Bennett, RR, Xia, Y, Ivanovska, IL, Liu, AJ, Greenberg, RA, and Discher, DE (2016). Nuclear constriction segregates mobile nuclear proteins away from chromatin. Mol Biol Cell 27, 4011–4020.

Jao, CY, and Salic, A (2008). Exploring RNA transcription and turnover in vivo by using click chemistry. Proc Natl Acad Sci U S A 105, 15779–15784.

Jiang, Y et al. (2020). Genome-wide analyses of chromatin interactions after the loss of Pol I, Pol II, and Pol III. Genome Biol 21, 158.

Kalukula, Y, Stephens, AD, Lammerding, J, and Gabriele, S (2022). Mechanics and functional consequences of nuclear deformations. Nat Rev Mol Cell Biol.

Khan, ZS, Santos, JM, and Hussain, F (2018). Aggressive prostate cancer cell nuclei have reduced stiffness. Biomicrofluidics 12, 014102.

Khatau, SB, Hale, CM, Stewart-Hutchinson, PJ, Patel, MS, Stewart, CL, Searson, PC, Hodzic, D, and Wirtz, D (2009). A perinuclear actin cap regulates nuclear shape. Proc Natl Acad Sci U S A 106, 19017–19022.

Kind, J, Pagie, L, Ortabozkoyun, H, Boyle, S, de Vries, SS, Janssen, H, Amendola, M, Nolen, LD, Bickmore, WA, and van Steensel, B (2013). Single-cell dynamics of genome-nuclear lamina interactions. Cell 153, 178–192.

Krause, M, Te Riet, J, and Wolf, K (2013). Probing the compressibility of tumor cell nuclei by combined atomic force-confocal microscopy. Phys Biol 10, 065002.

Lammerding, J, Fong, LG, Ji, JY, Reue, K, Stewart, CL, Young, SG, and Lee, RT (2006). Lamins A and C but not lamin B1 regulate nuclear mechanics. J Biol Chem 281, 25768–25780.

Le Berre, M, Aubertin, J, and Piel, M (2012). Fine control of nuclear confinement identifies a threshold deformation leading to lamina rupture and induction of specific genes. Integr Biol (Camb) 4, 1406–1414.

Lionetti, MC, Bonfanti, S, Fumagalli, MR, Budrikis, Z, Font-Clos, F, Costantini, G, Chepizhko, O, Zapperi, S, and La Porta, CAM (2020). Chromatin and cytoskeletal tethering determine nuclear morphology in progerin-expressing cells. Biophys J 118, 2319–2332.

Liu, K, Patteson, AE, Banigan, EJ, and Schwarz, JM (2021). Dynamic nuclear structure emerges from chromatin cross-links and motors. Phys Rev Lett 126, 158101.

Lu, C, Romo-Bucheli, D, Wang, X, Janowczyk, A, Ganesan, S, Gilmore, H, Rimm, D, and Madabhushi, A (2018). Nuclear shape and orientation features from H&E images predict survival in early-stage estrogen receptor-positive breast cancers. Lab Invest 98, 1438–1448.

Mahamid, J, Pfeffer, S, Schaffer, M, Villa, E, Danev, R, Cuellar, LK, Förster, F, Hyman, AA, Plitzko, JM, and Baumeister, W (2016). Visualizing the molecular sociology at the HeLa cell nuclear periphery. Science 351, 969–972.

Mistriotis, P et al. (2019). Confinement hinders motility by inducing RhoA-mediated nuclear influx, volume expansion, and blebbing. J Cell Biol 218, 4093–4111.

Nagashima, R et al. (2019). Single nucleosome imaging reveals loose genome chromatin networks via active RNA polymerase II. J Cell Biol 218, 1511–1530.

Nava, MM et al. (2020). Heterochromatin-driven nuclear softening protects the genome against mechanical stress-induced damage. Cell 181, 800–817.e22.

Nmezi, B et al. (2019). Concentric organization of A- and B-type lamins predicts their distinct roles in the spatial organization and stability of the nuclear lamina. Proc Natl Acad Sci U S A 116, 4307–4315.

Papanicolaou, GN, and Traut, HF (1997). The diagnostic value of vaginal smears in carcinoma of the uterus. 1941. Arch Pathol Lab Med 121, 211–224.

Patteson, AE, Vahabikashi, A, Pogoda, K, Adam, SA, Mandal, K, Kittisopikul, M, Sivagurunathan, S, Goldman, A, Goldman, RD, and Janmey, PA (2019). Vimentin protects cells against nuclear rupture and DNA damage during migration. J Cell Biol 218, 4079–4092.

Paulose, J, and Nelson, DR (2013). Buckling pathways in spherical shells with soft spots. Soft Matter 9, 8227.

Penfield, L, Wysolmerski, B, Mauro, M, Farhadifar, R, Martinez, MA, Biggs, R, Wu, H-Y, Broberg, C, Needleman, D, and Bahmanyar, S (2018). Dynein pulling forces counteract lamin-mediated nuclear stability during nuclear envelope repair. Mol Biol Cell 29, 852–868.

Pfeifer, CR et al. (2018). Constricted migration increases DNA damage and independently represses cell cycle. Mol Biol Cell 29, 1948–1962.

Pfeifer, CR, Tobin, MP, Cho, S, Vashisth, M, Dooling, LJ, Vazquez, LL, Ricci-De Lucca, EG, Simon, KT, and Discher, DE (2022). Gaussian curvature dilutes the nuclear lamina, favoring nuclear rupture, especially at high strain rate. Nucleus 13, 129–143.

Raab, M et al. (2016). ESCRT III repairs nuclear envelope ruptures during cell migration to limit DNA damage and cell death. Science 352, 359–362.

Radhakrishnan, A, Damodaran, K, Soylemezoglu, AC, Uhler, C, and Shivashankar, GV (2017). Machine learning for nuclear mechano-morphometric biomarkers in cancer diagnosis. Sci Rep 7.

Robijns, J, Molenberghs, F, Sieprath, T, Corne, TDJ, Verschuuren, M, and De Vos, WH (2016). In silico synchronization reveals regulators of nuclear ruptures in lamin A/C deficient model cells. Sci Rep 6, 30325.

Sapra, KT, Qin, Z, Dubrovsky-Gaupp, A, Aebi, U, Müller, DJ, Buehler, MJ, and Medalia, O (2020). Nonlinear mechanics of lamin filaments and the meshwork topology build an emergent nuclear lamina. Nat Commun 11, 6205.

Schreiner, SM, Koo, PK, Zhao, Y, Mochrie, SGJ, and King, MC (2015). The tethering of chromatin to the nuclear envelope supports nuclear mechanics. Nat Commun 6, 7159.

Senigagliesi, B et al. (2019). The High Mobility Group A1 (HMGA1) chromatin architectural factor modulates nuclear stiffness in breast cancer cells. Int J Mol Sci 20, 2733.

Shaban, HA, Barth, R, and Bystricky, K (2018). Formation of correlated chromatin domains at nanoscale dynamic resolution during transcription. Nucleic Acids Res 46, e77–e77.

Shaban, HA, Barth, R, Recoules, L, and Bystricky, K (2020). Hi-D: nanoscale mapping of nuclear dynamics in single living cells. Genome Biol 21, 95.

Shah, P, Hobson, CM, Cheng, S, Colville, MJ, Paszek, MJ, Superfine, R, and Lammerding, J (2021). Nuclear deformation causes DNA damage by increasing replication stress. Curr Biol 31, 753–765.e6.

Shimamoto, Y, Tamura, S, Masumoto, H, and Maeshima, K (2017). Nucleosome-nucleosome interactions via histone tails and linker DNA regulate nuclear rigidity. Mol Biol Cell 28, 1580–1589.

Shimi, T et al. (2008). The A- and B-type nuclear lamin networks: microdomains involved in chromatin organization and transcription. Genes Dev 22, 3409–3421.

Shimi, T, Kittisopikul, M, Tran, J, Goldman, AE, Adam, SA, Zheng, Y, Jaqaman, K, and Goldman, RD (2015). Structural organization of nuclear lamins A, C, B1, and B2 revealed by superresolution microscopy. Mol Biol Cell 26, 4075–4086.

Srivastava, N, Nader, GP de F, Williart, A, Rollin, R, Cuvelier, D, Lomakin, A, and Piel, M (2021). Nuclear fragility, blaming the blebs. Curr Opin Cell Biol 70, 100–108.

Stephens, AD et al. (2019a). Physicochemical mechanotransduction alters nuclear shape and mechanics via heterochromatin formation. Mol Biol Cell 30, 2320–2330.

Stephens, AD (2020). Chromatin rigidity provides mechanical and genome protection. Mutat Res 821, 111712.

Stephens, AD, Banigan, EJ, Adam, SA, Goldman, RD, and Marko, JF (2017). Chromatin and lamin A determine two different mechanical response regimes of the cell nucleus. Mol Biol Cell 28, 1984–1996.

Stephens, AD, Banigan, EJ, and Marko, JF (2019b). Chromatin’s physical properties shape the nucleus and its functions. Curr Opin Cell Biol 58, 76–84.

Stephens, AD, Liu, PZ, Banigan, EJ, Almassalha, LM, Backman, V, Adam, SA, Goldman, RD, and Marko, JF (2018). Chromatin histone modifications and rigidity affect nuclear morphology independent of lamins. Mol Biol Cell 29, 220–233.

Strickfaden, H, Tolsma, TO, Sharma, A, Underhill, DA, Hansen, JC, and Hendzel, MJ (2020). Condensed chromatin behaves like a solid on the mesoscale in vitro and in living cells. Cell 183, 1772–1784.e13.

Strom, AR et al. (2021). HP1α is a chromatin crosslinker that controls nuclear and mitotic chromosome mechanics. Elife 10.

Turgay, Y, Eibauer, M, Goldman, AE, Shimi, T, Khayat, M, Ben-Harush, K, Dubrovsky-Gaupp, A, Sapra, KT, Goldman, RD, and Medalia, O (2017). The molecular architecture of lamins in somatic cells. Nature 543, 261–264.

Vahabikashi, A et al. (2022). Nuclear lamin isoforms differentially contribute to LINC complex-dependent nucleocytoskeletal coupling and whole-cell mechanics. Proc Natl Acad Sci U S A 119, e2121816119.

Vargas, JD, Hatch, EM, Anderson, DJ, and Hetzer, MW (2012). Transient nuclear envelope rupturing during interphase in human cancer cells. Nucleus 3, 88–100.

Wang, P et al. (2018). WDR5 modulates cell motility and morphology and controls nuclear changes induced by a 3D environment. Proc Natl Acad Sci U S A 115, 8581–8586.

Williams, JF, Surovtsev, IV, Schreiner, SM, Nguyen, H, Hu, Y, Mochrie, SGJ, and King, MC (2020). Phase separation enables heterochromatin domains to do mechanical work, bioRxiv.

Wren, NS, Zhong, Z, Schwartz, RS, and Dahl, KN (2012). Modeling nuclear blebs in a nucleoskeleton of independent filament networks. Cell Mol Bioeng 5, 73–81.

Xia, Y et al. (2018). Nuclear rupture at sites of high curvature compromises retention of DNA repair factors. J Cell Biol 217, 3796–3808.

Yong, EH, Nelson, DR, and Mahadevan, L (2013). Elastic platonic shells. Phys Rev Lett 111, 177801.

Young, AM, Gunn, AL, and Hatch, EM (2020). BAF facilitates interphase nuclear membrane repair through recruitment of nuclear transmembrane proteins. Mol Biol Cell 31, 1551–1560.

Zhang, Q, Tamashunas, AC, Agrawal, A, Torbati, M, Katiyar, A, Dickinson, RB, Lammerding, J, and Lele, TP (2019). Local, transient tensile stress on the nuclear membrane causes membrane rupture. Mol Biol Cell 30, 899–906.

Zhang, S et al. (2021). RNA polymerase II is required for spatial chromatin reorganization following exit from mitosis. Sci Adv 7, eabg8205.

Zhang, S, Uebelmesser, N, Barbieri, M, and Papantonis, A (2022). Enhancer-promoter contact formation requires RNAPII and antagonizes loop extrusion.

Zidovska, A, Weitz, DA, and Mitchison, TJ (2013). Micron-scale coherence in interphase chromatin dynamics. Proc Natl Acad Sci U S A 110, 15555–15560.

